# Meta-omics-aided isolation of an elusive anaerobic arsenic-methylating soil bacterium

**DOI:** 10.1101/2022.01.25.477449

**Authors:** Karen Viacava, Jiangtao Qiao, Andrew Janowczyk, Suresh Poudel, Nicolas Jacquemin, Karin Lederballe Meibom, Him K. Shrestha, Matthew C. Reid, Robert L. Hettich, Rizlan Bernier-Latmani

**Author notes:** Corresponding author. E-mail address. Postal address EPFL ENAC IIE EML CH A1 375 (Bâtiment CH) Station 6 CH-1015 Lausanne Switzerland.

## Abstract

Soil microbiomes harbor unparalleled functional and phylogenetic diversity and are sources of novel metabolisms. However, extracting isolates with a targeted function from complex microbiomes is not straightforward, particularly if the associated phenotype does not lend itself to high-throughput screening. Here, we tackle the methylation of arsenic (As) in anoxic soils. By analogy to mercury, As methylation was proposed to be catalyzed by sulfate-reducing bacteria. However, to date, there are no anaerobic isolates capable of As methylation, whether sulfate-reducing or otherwise. The isolation of such a microorganism has been thwarted by the fact that the anaerobic bacteria harboring a functional arsenite S-adenosylmethionine methyltransferase (ArsM) tested to date did not methylate As in pure culture. Additionally, fortuitous As methylation can result from the release of non-specific methyltransferases upon lysis. Thus, we combined metagenomics, metatranscriptomics, and metaproteomics to identify the microorganisms actively methylating As in anoxic soil-derived microbial cultures. Based on the metagenome-assembled genomes of microorganisms expressing ArsM, we isolated *Paraclostridium* sp. strain EML, which was confirmed to actively methylate As anaerobically. This work is an example of the application of meta-omics to the isolation of elusive microorganisms.

## Introduction

Soil microbiomes represent a rich source of novel metabolisms and taxa. However, isolating microorganisms to study specific functions from these microbiomes can be challenging, and even more so in cases for which the phenotype is not identifiable with high-throughput methods. An example of challenging microorganisms to identify are anaerobic As-methylating strains. Arsenic is a toxic metalloid that is naturally-occurring, extensively distributed in the environment, and the substrate of numerous microbial transformations [1]. One such transformation is As methylation, catalyzed by arsenite (As(III)) S-adenosylmethionine methyltransferase (ArsM) [2], which entails the binding of one to three methyl group(s) to the As atom. Arsenic methylation occurs in flooded rice paddy soils, resulting in the accumulation of methylated As in rice grains [3]. The bioaccumulation of methylated As in rice grains is considerably more efficient than that of inorganic As species [4, 5].

The gene encoding ArsM (*arsM*) has been identified in phylogenetically diverse soil microorganisms [6–9]. The production of toxic trivalent monomethylated As (MMAs(III)) by anaerobic prokaryotes has been proposed as a microbial warfare strategy, to inhibit microbial competitors with what amounts to an arsenic-containing antibiotic [10]. If that is the case, it is conceivable that As methylation may not occur in pure cultures but only in microbial communities, triggered by metabolites produced by the microbiome. Alternatively, *arsM*-harboring microorganisms that express As(III) efflux pump(s), the major pathway of As resistance within bacteria [11], may not methylate As due to the efficient removal of As(III) from the cytoplasm, which is the location of ArsM [12, 13]. Either occurrence would render the isolation of pure cultures of As-methylating anaerobes very challenging using standard approaches. The latter hypothesis is supported by recent work showing the lack of As methylation by anaerobic pure cultures harboring functional ArsM enzymes [13].

An additional complexity is the evidence for the fortuitous methylation of As upon cell lysis and release of methyltransferases. This occurrence was suggested by considering the methanogen *Methanosarcina mazei* for which As methylation was initiated only when cell viability decreased [13]. Thus, presence of As methylation for cultures incubated beyond the exponential phase may only be an experimental artefact. Finally, the detection of methylated As requires relatively complex analytical tools (high pressure liquid chromatography coupled to inductively-coupled plasma mass spectrometry, HPLC-ICP-MS) that do not lend themselves readily to high-throughput screening of a large number of colonies. As a result of these challenges, there are no anaerobic microorganisms known to actively methylate As despite many efforts to identify them. In one instance, researchers had identified a Gram-positive sulfate-reducing bacterium (SRB) [14] that was reported to methylate As but this isolate is no longer available, precluding further investigation.

Thus, this study aimed to conclusively identify an active anaerobic As methylator in soil-derived microbial cultures using a multi-omics approach. The experimental strategy was to build Metagenome-Assembled Genomes (MAGs) from metagenomic data of the microbiome and to identify the subset of MAGs harboring the gene *arsM* that also expressed *arsM* (metatranscriptomics) and/or ArsM (metaproteomics). Based on the genetic information from the target MAG, an informed isolation strategy was devised that allowed the recovery of a pure culture later confirmed to be a novel anoxic As-methylating strain.

## Materials and methods

### Rice paddy soil microbiomes

The soil-derived cultures consist of two anaerobic cultures derived from a Vietnamese rice paddy soil and introduced in Reid *et al*. [15]. The first soil-derived microbiome was grown in ¼ strength tryptic soy broth (TSB) medium (7.5 g l^-1^ TSB), used previously to enrich arsenic-methylating microbes from a lake sediment [16], and will be referred to as the TSB culture. The medium from the second soil-derived microbiome, in addition to ¼ strength TSB, included electron acceptors and two additional carbon sources to simultaneously allow the growth of nitrate-, iron-, and sulfate-reducers, as well as microbes with fermentative and methanogenic metabolisms (EA medium: 5 mM NaNO_3_, 5 mM Na_2_SO_4_, 5 mM ferric citrate, 0.2 g l^-1^ yeast extract (Oxoid, Hampshire, UK) and 1 g l^-1^ cellobiose, pH 7). This enrichment will be referred to as the EA culture. Both media were boiled, cooled down under 100% N_2_ gas and 50 ml of medium were dispensed into 100-ml serum bottles. The bottle headspace was flushed with 100% N_2_ gas prior to autoclaving. All culture manipulations were carried out using thoroughly N_2_-flushed syringes and needles. Cultures were grown at 30°C. Growth was quantified using optical density at 600 nm (OD_600_).

### Arsenic methylation assays

Pre-cultures from each culture were started from −80°C glycerol stocks. The EA culture started from the glycerol stock was transferred only after a dark precipitate, presumably iron sulfide (suggesting potentially active sulfate reduction), was formed. The first experimental set-up consisted of the inoculation of bottles containing medium amended with As(III) as NaAsO_2_ (+As condition) or unamended (no-As control). Cell pellets were sampled during the stationary phase for DNA sequencing and proteome characterization and at the mid-exponential growth phase for RNA sequencing (see Figures S1, S2 and S3 Supplementary Information (SI) for precise times). In a second experimental set-up, cultures were grown in medium without As(III) and, at the mid-exponential growth phase, As(III) was added. Cell pellets were sampled before (no-As control) and 30 min after As spiking (+As condition) and were used for a second transcriptomic analysis only. All experiments were performed in biological triplicates. Sampling for soluble As species, determination of As speciation and total As concentration were performed as described in [13] using an Agilent 8900 ICP-QQQ instrument coupled to an HPLC 1260 Infinity II (Agilent Technologies, CA, USA). Instrument settings in Table S1. The DNA and RNA sequencing, metaproteome characterization, metagenomic, metatranscriptomic, and metaproteomic analyses are described in SI.

### *Isolation of* As-methylating microorganism

The isolation of the anaerobic *arsM*-expressing microorganism was conducted by using serial dilution agar plate method in an anaerobic chamber (Coy Laboratories, Grass Lake, MI, USA) containing 90% N_2_:10% H_2_ with less than 5 ppm of O_2_. Briefly, 1 ml of cell suspension of the EA culture was serially diluted in a 10-fold series (10^-1^ to 10^-5^ dilutions) using sterile Reinforced Clostridial Broth (RCB) (Oxoid Ltd., Basingstoke, UK). Consecutively, 100 μl of EA cell suspension from each dilution was spread uniformly over the surface of Reinforced Clostridial agar (RCA) (Oxoid Ltd., Basingstoke, UK). The inoculated RCA plates were incubated at 30°C for 24 hours. The single colonies were transferred with sterile toothpicks to Tryptose-sulfite-cycloserine (TSC) agar (Merck, Darmstadt), containing 5 g l^-1^ sucrose, 0.04 g l^-1^ bromocresol purple (Sigma-Aldrich), and 0.4 g l^-1^ de-hydrated D-cycloserine (Sigma-Aldrich), and grown at 30°C for 24 hours.

Colony PCR was performed on black colonies picked from purple TSC agar using the designed specific primers arsM-9F: 5’-TCTAATCTAAGTTGTTGTGGGGAAG-3’ and arsM-9R: 5’-TGATATAGATAACCTACCTCCGCC-3’, generating a 500 bp amplicon of the *arsM* gene from MAG 9 from the EA culture (Table 1-A) (protein id k119_30669_28, Table S2). Before direct colony PCR, the black colonies were first picked, diluted into 15 μl lysis buffer (0.1% triton X-100 + TE buffer) and boiled at 95°C for 10 min, to release DNA, and then centrifuged (13,000 g, 10 min) to spin down cell debris. The supernatant of the lysate was used as the DNA template for colony PCR.

**Table 1.**
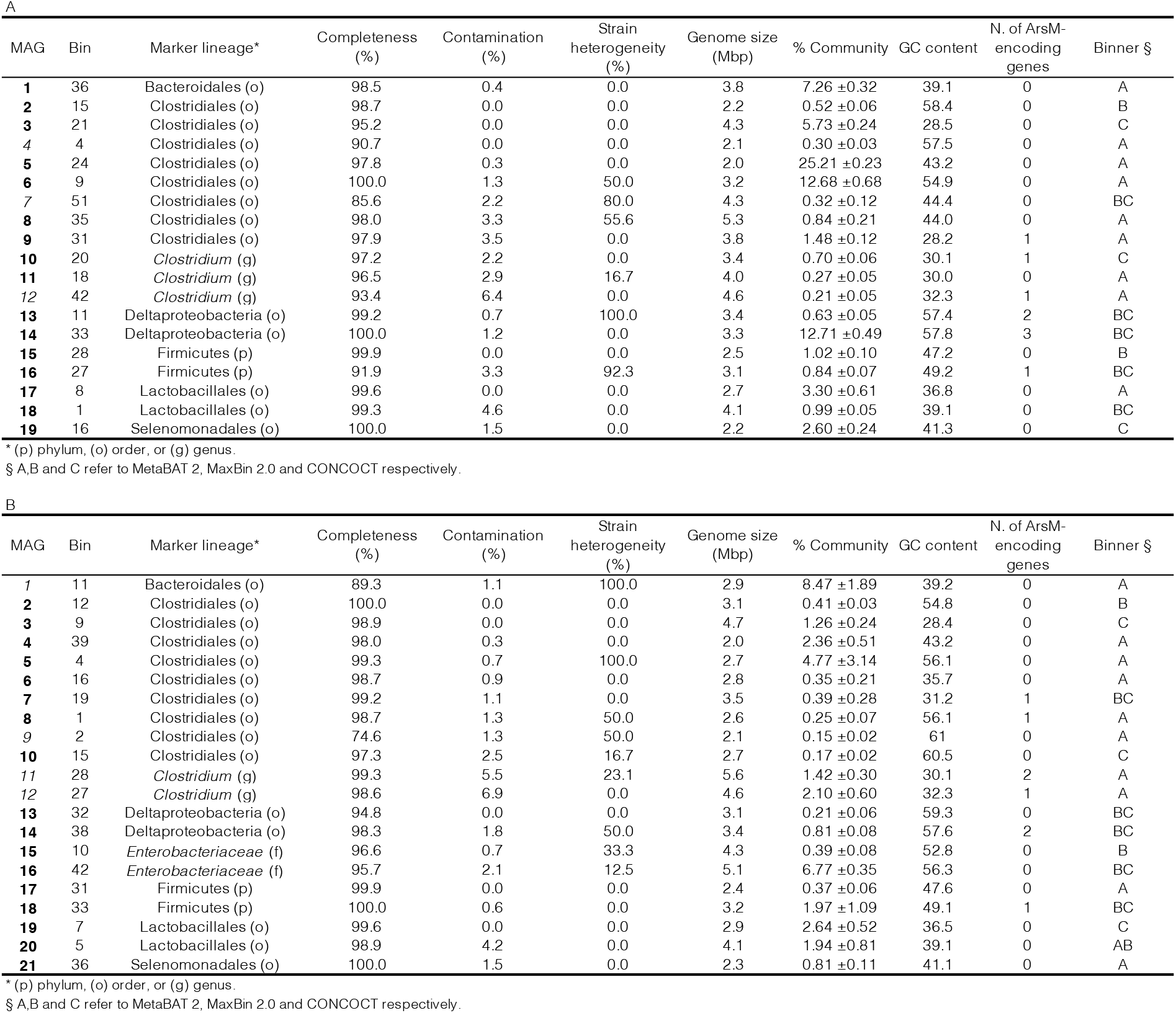
Metagenome assembled genomes (MAGs) from EA (upper Table A) and TSB (lower Table B) cultures in the +As condition. Marker lineage: taxonomic rank set by CheckM. Completeness and contamination %: estimated completeness and contamination of genome as determined by CheckM from the presence/absence of single-copy marker genes and the expected colocalization of these genes. Strain heterogeneity: index between 0 and 100 where a value of 0 means no strain heterogeneity, high values suggest the majority of reported contamination is from closely related organisms (i.e., potentially the same species) and low values suggest the majority of contamination is from phylogenetically diverse sources. % of binned proteins assigned to MAG: number of protein-coding genes assigned to the MAG divided by the total number of protein-coding genes binned. % Community: sum of the number of reads mapped to the contigs in each MAG divided by the total number of reads mapped to all contigs including the unbinned contigs, and normalized to MAG size, assuming an average genome size for all unbinned populations. High-quality MAGs are denoted by bolded numbers, good-quality MAGs by italicized numbers.

| A |  |  |  |  |  |  |  |  |  |  |
| --- | --- | --- | --- | --- | --- | --- | --- | --- | --- | --- |
| MAG | Bin | Marker lineage* | Completeness (%) | Contamination (%) | Strain heterogeneity (%) | Genome size (Mbp) | % Community | GC content | N. of ArsM-encoding genes | Binner § |
| **1** | 36 | Bacteroidales (o) | 98.5 | 0.4 | 0.0 | 3.8 | 7.26 ±0.32 | 39.1 | 0 | A |
| **2** | 15 | Clostridiales (o) | 98.7 | 0.0 | 0.0 | 2.2 | 0.52 ±0.06 | 58.4 | 0 | B |
| **3** | 21 | Clostridiales (o) | 95.2 | 0.0 | 0.0 | 4.3 | 5.73 ±0.24 | 28.5 | 0 | C |
| **4** | 4 | Clostridiales (o) | 90.7 | 0.0 | 0.0 | 2.1 | 0.30 ±0.03 | 57.5 | 0 | A |
| **5** | 24 | Clostridiales (o) | 97.8 | 0.3 | 0.0 | 2.0 | 25.21 ±0.23 | 43.2 | 0 | A |
| **6** | 9 | Clostridiales (o) | 100.0 | 1.3 | 50.0 | 3.2 | 12.68 ±0.68 | 54.9 | 0 | A |
| **7** | 51 | Clostridiales (o) | 85.6 | 2.2 | 80.0 | 4.3 | 0.32 ±0.12 | 44.4 | 0 | BC |
| **8** | 35 | Clostridiales (o) | 98.0 | 3.3 | 55.6 | 5.3 | 0.84 ±0.21 | 44.0 | 0 | A |
| **9** | 31 | Clostridiales (o) | 97.9 | 3.5 | 0.0 | 3.8 | 1.48 ±0.12 | 28.2 | 1 | A |
| **10** | 20 | *Clostridium* (g) | 97.2 | 2.2 | 0.0 | 3.4 | 0.70 ±0.06 | 30.1 | 1 | C |
| **11** | 18 | *Clostridium* (g) | 96.5 | 2.9 | 16.7 | 4.0 | 0.27 ±0.05 | 30.0 | 0 | A |
| **12** | 42 | *Clostridium* (g) | 93.4 | 6.4 | 0.0 | 4.6 | 0.21 ±0.05 | 32.3 | 1 | A |
| **13** | 11 | Deltaproteobacteria (o) | 99.2 | 0.7 | 100.0 | 3.4 | 0.63 ±0.05 | 57.4 | 2 | BC |
| **14** | 33 | Deltaproteobacteria (o) | 100.0 | 1.2 | 0.0 | 3.3 | 12.71 ±0.49 | 57.8 | 3 | BC |
| **15** | 28 | Firmicutes (p) | 99.9 | 0.0 | 0.0 | 2.5 | 1.02 ±0.10 | 47.2 | 0 | B |
| **16** | 27 | Firmicutes (p) | 91.9 | 3.3 | 92.3 | 3.1 | 0.84 ±0.07 | 49.2 | 1 | BC |
| **17** | 8 | Lactobacillales (o) | 99.6 | 0.0 | 0.0 | 2.7 | 3.30 ±0.61 | 36.8 | 0 | A |
| **18** | 1 | Lactobacillales (o) | 99.3 | 4.6 | 0.0 | 4.1 | 0.99 ±0.05 | 39.1 | 0 | BC |
| **19** | 16 | Selenomonadales (o) | 100.0 | 1.5 | 0.0 | 2.2 | 2.60 ±0.24 | 41.3 | 0 | C |

\* (p) phylum, (o) order, or (g) genus.
§ A, B and C refer to MetaBAT 2, MaxBin 2.0 and CONCOCT respectively.

| B |  |  |  |  |  |  |  |  |  |  |
| --- | --- | --- | --- | --- | --- | --- | --- | --- | --- | --- |
| MAG | Bin | Marker lineage* | Completeness (%) | Contamination (%) | Strain heterogeneity (%) | Genome size (Mbp) | % Community | GC content | N. of ArsM-encoding genes | Binner § |
| **1** | 11 | Bacteroidales (o) | 89.3 | 1.1 | 100.0 | 2.9 | 8.47 ±1.89 | 39.2 | 0 | A |
| **2** | 12 | Clostridiales (o) | 100.0 | 0.0 | 0.0 | 3.1 | 0.41 ±0.03 | 54.8 | 0 | B |
| **3** | 9 | Clostridiales (o) | 98.9 | 0.0 | 0.0 | 4.7 | 1.26 ±0.24 | 28.4 | 0 | C |
| **4** | 39 | Clostridiales (o) | 98.0 | 0.3 | 0.0 | 2.0 | 2.36 ±0.51 | 43.2 | 0 | A |
| **5** | 4 | Clostridiales (o) | 99.3 | 0.7 | 100.0 | 2.7 | 4.77 ±3.14 | 56.1 | 0 | A |
| **6** | 16 | Clostridiales (o) | 98.7 | 0.9 | 0.0 | 2.8 | 0.35 ±0.21 | 35.7 | 0 | A |
| **7** | 19 | Clostridiales (o) | 99.2 | 1.1 | 0.0 | 3.5 | 0.39 ±0.28 | 31.2 | 1 | BC |
| **8** | 1 | Clostridiales (o) | 98.7 | 1.3 | 50.0 | 2.6 | 0.25 ±0.07 | 56.1 | 1 | A |
| **9** | 2 | Clostridiales (o) | 74.6 | 1.3 | 50.0 | 2.1 | 0.15 ±0.02 | 61 | 0 | A |
| **10** | 15 | Clostridiales (o) | 97.3 | 2.5 | 16.7 | 2.7 | 0.17 ±0.02 | 60.5 | 0 | C |
| **11** | 28 | *Clostridium* (g) | 99.3 | 5.5 | 23.1 | 5.6 | 1.42 ±0.30 | 30.1 | 2 | A |
| **12** | 27 | *Clostridium* (g) | 98.6 | 6.9 | 0.0 | 4.6 | 2.10 ±0.60 | 32.3 | 1 | A |
| **13** | 32 | Deltaproteobacteria (o) | 94.8 | 0.0 | 0.0 | 3.1 | 0.21 ±0.06 | 59.3 | 0 | BC |
| **14** | 38 | Deltaproteobacteria (o) | 98.3 | 1.8 | 50.0 | 3.4 | 0.81 ±0.08 | 57.6 | 2 | BC |
| **15** | 10 | *Enterobacteriaceae* (f) | 96.6 | 0.7 | 33.3 | 4.3 | 0.39 ±0.08 | 52.8 | 0 | B |
| **16** | 42 | *Enterobacteriaceae* (f) | 95.7 | 2.1 | 12.5 | 5.1 | 6.77 ±0.35 | 56.3 | 0 | BC |
| **17** | 31 | Firmicutes (p) | 99.9 | 0.0 | 0.0 | 2.4 | 0.37 ±0.06 | 47.6 | 0 | A |
| **18** | 33 | Firmicutes (p) | 100.0 | 0.6 | 0.0 | 3.2 | 1.97 ±1.09 | 49.1 | 1 | BC |
| **19** | 7 | Lactobacillales (o) | 99.6 | 0.0 | 0.0 | 2.9 | 2.64 ±0.52 | 36.5 | 0 | C |
| **20** | 5 | Lactobacillales (o) | 98.9 | 4.2 | 0.0 | 4.1 | 1.94 ±0.81 | 39.1 | 0 | AB |
| **21** | 36 | Selenomonadales (o) | 100.0 | 1.5 | 0.0 | 2.3 | 0.81 ±0.11 | 41.1 | 0 | A |

\* (p) phylum, (o) order, or (g) genus.
§ A, B and C refer to MetaBAT 2, MaxBin 2.0 and CONCOCT respectively.

Each colony PCR consisted of a 25-μl reaction containing: 12.5 μl 2x GoTaq Green Master mix (Promega, UK), 0.5 μl of the reverse and forward primers (10 μM each), 1.5 μl of lysate supernatant as the DNA template, 0.25 μl of 20 mg ml^-1^ bovine serum albumin (BSA) (Sigma-Aldrich), and 10.75 μl sterile DNase RNase Free water. The thermocycling program consisted of an initial denaturation at 95°C for 5 min, 30 cycles of denaturation at 95°C for 40s, annealing at 53°C for 40 sec, and extension at 72°C for 40s, and a final extension at 72°C for 10 min. After purification of the PCR product with Wizard® SV Gel and PCR Clean-Up System (Promega, UK), the *arsM* amplicon was sequenced at Microsynth (Balgach, Switzerland). The amplification of full-length 16S rRNA gene was performed using the primers 27F [17] and 1542R [18]. The As methylation assay of the isolate is described in SI.

## Results

### Arsenic methylation by soil-derived microbiomes

As described in materials and methods, two experimental set-ups were used to probe the As-methylating cultures. The first set-up compared the soil-derived microbial communities grown in the presence or absence of As(III). Samples for the metagenome, metaproteome and one of the metatranscriptomes (labeled metatranscriptome G for ‘growth in the presence of As’) were obtained from this set-up (Figures S1, S2 and S3). In the second set-up, the intention was to capture the short-term response of the community to As(III). Thus, the cultures were grown to mid-exponential phase in the absence of As, sampled, amended with As(III), and sampled a second time after 30 minutes. Samples obtained from this set-up were used for the second metatranscriptome (labeled metatranscriptome R for ‘response to arsenic addition’) (Figures S1-A and S2-A).

Both cultures exhibited As methylation, reaching an efficiency of transformation of the initial As(III) of 27.7% and 19.5% for the EA and TSB cultures, respectively (Figures S1 and S2 and SI). The TSB and EA microbiomes compare favorably to previous studies of anaerobic enrichments [16] and single-strain cultures [13, 14, 19], and represent the most efficient anoxic As-methylating microbial cultures reported.

### Microbiome composition

The taxonomic classification of small subunit (SSU) 16S rRNA sequences show that, although eukaryotic DNA was also identified, the main fraction of the communities is bacterial (>89.0 ±0.8% for EA cultures and >98.5 ±0.3% for TSB cultures, relative abundance) and is distributed amongst eight operational taxonomic units (OTUs) at the order level: Acidaminococcales, Bacillales, Bacteroidales, Clostridiales, Desulfovibrionales, Enterobacterales, Lactobacillales and Selenomonadales (Figure 1 and Tables S3-S6). Statistically significant changes in the OTUs relative abundances, +As condition versus no-As control, are described in SI.

**Figure 1.**
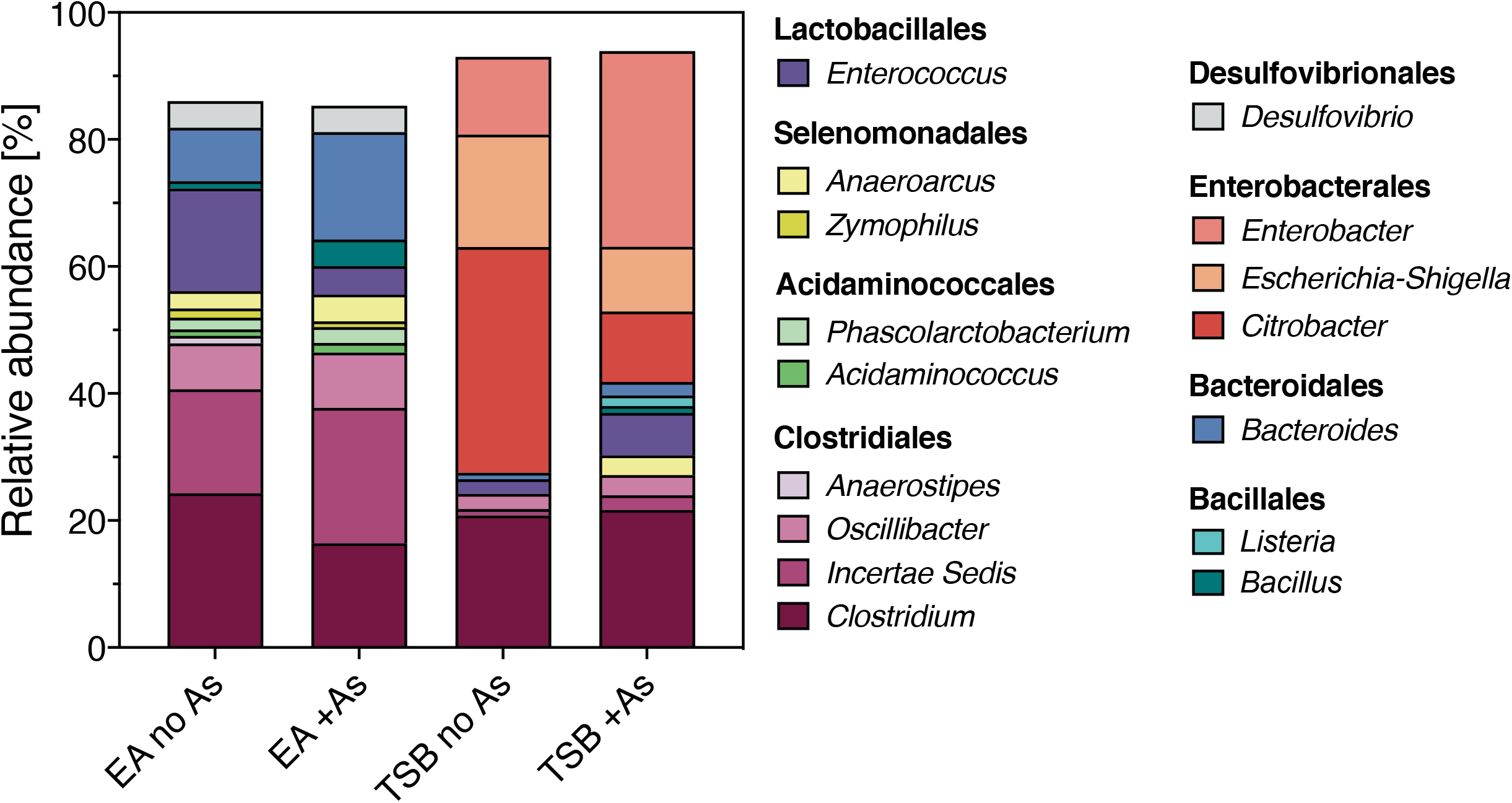
Operational taxonomic units (OTUs) at order and genus level (with > 1% relative abundance at genus level) identified from SSU 16S rRNA sequences from soil-derived cultures. Abbreviations: EA no As: EA culture no-As control EA +As: EA culture +As condition, TSB no As: TSB culture no-As control TSB +As: TSB culture +As condition. OTUs at the order level are indicated in bold in the legend. Plotted values are the average relative abundance and together with SD values and student t-test results are available in SI Tables S14 and S15.

The read-assembled contiguous sequences (contigs) from the four metagenomes, EA (+As, no-As control) and TSB (+As, no-As control), were clustered separately into bins. High-quality (≥95% completeness and ≤5% contamination) and good-quality (≥74% completeness and ≤10% contamination) bins were designated as MAGs. For the +As condition, the parsing process led to a total of 40 MAGs: 19 in EA culture (Table 1-A), and 21 in TSB culture (Table 1-B). Additionally, matching bins for the +As condition, based on the MAG groups, were found in the bins from the no-As control cultures (Tables S7 and S8). Only three of the 40 MAGs in the +As condition were left unpaired (EA MAGs 7 and 12 and TSB MAG 10).

For each MAG group, a lineage was assigned by CheckM, based on lineage-specific marker genes [20]. The MAGs identified belonged mainly to the phylum Firmicutes (orders Clostridiales, Selenomonadales and Lactobacillales, and the genus *Clostridium*). Proteobacteria MAGs included the *Enterobacteriaceae* family and the Deltaproteobacteria class. Finally, in each metagenome, one MAG from the order Bacteroidales was present.

Fifteen MAGs presented strain heterogeneity, an index of the phylogenetic relatedness of binned contigs based on the amino acid identity of the encoded proteins. For ten MAGs, the value is >50%, suggesting some phylogenetic relation of the contaminating strains. Five MAGs had heterogeneity values <33.33%, suggesting contamination with microorganisms that are not closely related. In the remaining 25 MAGs, the strain heterogeneity is 0%, i.e., no strain heterogeneity or no contamination (Tables S7 and S8).

Ultimately, to evaluate the relatedness of the EA and TSB microbial communities, matching bins between the EA and TSB +As condition MAGs were found by pairwise comparison of the predicted genomes. Of the 40 MAGs identified across EA and TSB cultures, 22 (corresponding to 11 pairs) exhibited similarities >98.47% (Table S9), including a Deltaproteobacteria, a *Clostridium*, three Clostridiales, a Bacteroidales, two Lactobacillales, a Selenomonadales, and two Firmicutes. Thus, approximately half of the MAGs of the EA culture are also present in the TSB culture and vice-versa.

Changes in the relative abundance of MAGs (no-As control vs. +As condition), as well as the presence, transcription and translation of genes encoding key enzymes from major metabolic pathways from each MAG in the +As condition cultures are included in SI.

### Arsenic resistance genes

The metagenomic libraries from the +As condition of the EA and TSB cultures were mined for arsenic resistance (*ars*) genes and their encoded proteins (pipeline described in SI). A gene was considered to be present in the culture if DNA reads represented >5 TPM-DNA (‘transcripts per million’ (TPM), referred to as TPM-DNA when used for gene abundance, see SI), considered to be transcribed if >5 TPM-RNA (referred as TPM-RNA when employed for transcript abundance) were detected, and as translated if protein abundance could be calculated from the detected peptides in at least two of the three biological replicates. Additionally, increased expression in the RNA and protein in the +As condition relative to the no-As condition, was considered when the absolute log_2_ fold change was ≥1 (i.e., 0.5 ≥ fold change ≥2) and the adjusted q-value ≤0.05 (refer to SI for pipeline) (Figure 2). A total of 309 and 282 genes were annotated as *ars* genes in the EA and TSB +As metagenomic libraries, respectively (Tables S10 and S11). Of those, 255 and 226 were considered correctly annotated as *ars* genes based on BLAST® and HMMER (refer to SI for pipeline), and 225 and 147 had above-threshold DNA abundances, respectively (Figure 2). Individual values of transcript and protein abundance in the +As condition and the no-As control and their abundance values in the +As condition relative to the no-As control for each MAG group, biological replicate and *ars* proteins from the EA and TSB cultures are available in Tables S2 and S12, respectively.

**Figure 2.**
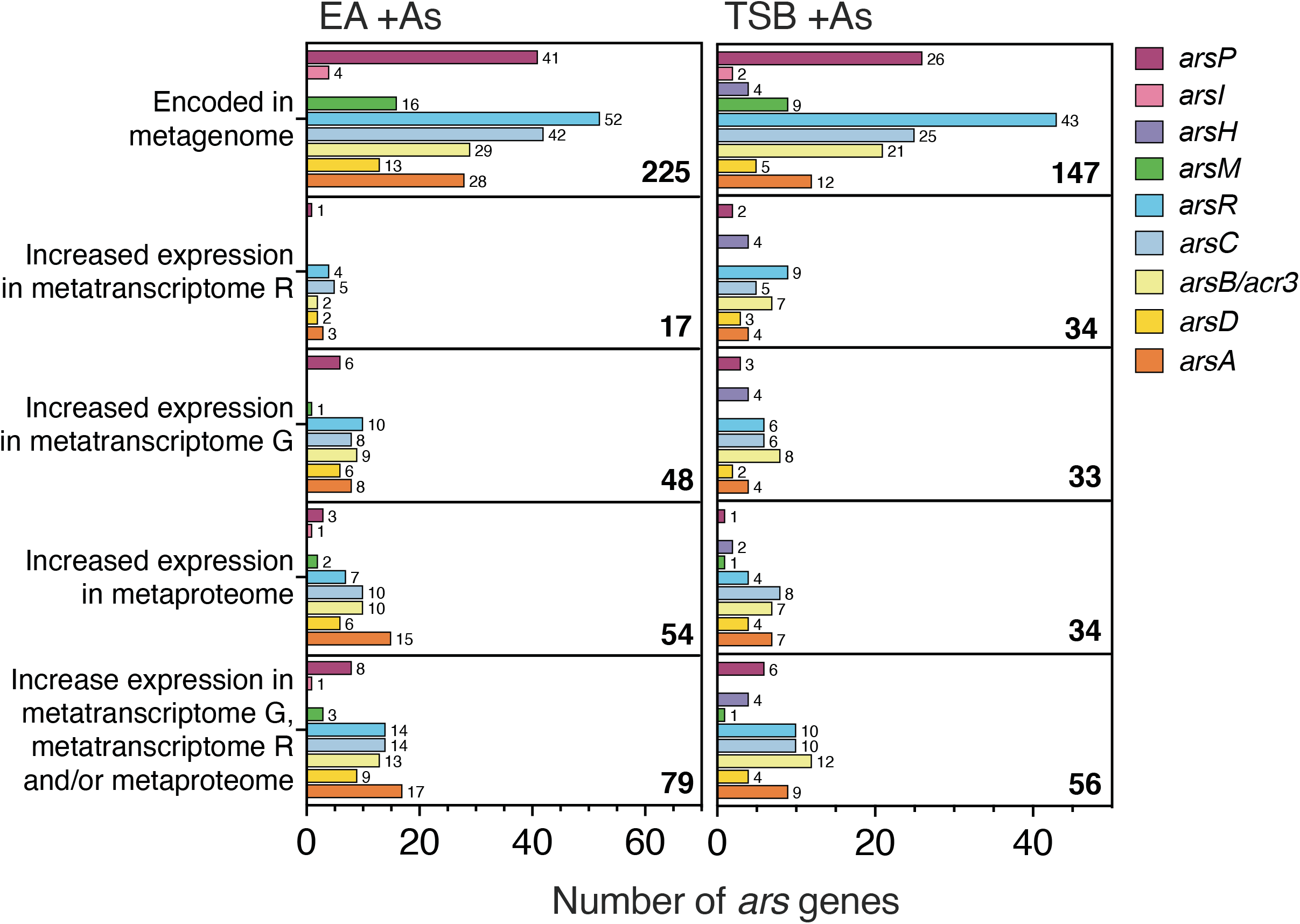
Number of *ars* genes, encoded in the +As condition cultures and with increased expression in metatranscriptomes/metaproteome relative to no-As controls. Number of *ars* genes encoded in metagenomes and with increased expression in metatranscriptomes, R or G, or metaproteomes and the non-redundant overlap between genes with increased expression in metatranscriptomes and/or metaproteomes from +As condition EA (left panels) and TSB (right panels) cultures. Bar length and numbers on the right side of the bars correspond to the number of genes per *ars* gene category. Bold numbers on the lower left corner of each panel correspond to the sum of all *ars* genes per category.

The *ars* genes encode proteins involved in the detoxification of As oxyanions: *arsB* and *acr3* encoding As(III)-efflux systems; *arsA* encoding the ATPase energizing the efflux of As(III) and mostly found alongside the gene for a As(III) chaperone and weak *ars* operon repressor (*arsD*) [21]; *arsC1* and *arsC2* encoding As(V) reductases coupling As reduction to the oxidation of glutaredoxin or thioredoxin, respectively; and *arsR* genes encoding As(III)-regulated repressors (ArsR1, ArsR2, and ArsR3) which are distinguishable based on the location of the As(III)-binding cysteine residues [22–24].

The most common *ars* genes in EA and TSB culture metagenomes were *arsR, arsC*, and *arsP* (Figure 2). The first two genes are part of the canonical *ars* operon *arsRBC* [25], whilst *arsP*, encoding a more recently discovered membrane transporter, has been found to be widely distributed in bacterial genomes [11]. Most of the surveyed *arsP* genes, 57% in EA and 50% in TSB, are encoded in putative *ars* operons, represented by *ars* genes contiguously encoded in the same contig (Tables S2 and S12), supporting their As-related function and correct annotation. Unsurprisingly, the genes responsible for As(III) efflux (*arsB, acr3*, and *arsA*), typically found in organisms living in reducing environments, were amongst the most common *ars* genes along with *arsC* [8, 26]. Finally, *arsM* and the two genes, *arsI* and *arsH*, encoding oxygen-dependent MMAs(III)-resistance mechanisms, were the least recurrent genes in the metagenomes. In soils, *arsM* has previously been observed to be less common in comparison to other surveyed *ars* genes [6, 8, 27]. The results of gene and protein relative expression vs. the no-As control of the *ars* genes involved in the metabolism of inorganic As in the MAGs can be found in SI.

It is striking to note that while there are a large number of *ars* genes in the metagenome, a small proportion is expressed (whether as mRNA transcripts or as proteins) in the presence of As when compared to the no As control (Figure 2). This contrast is particularly evident for the gene responsible for As methylation, *arsM*.

### Arsenic-methylating MAGs

The aim of the present study was to identify the microorganisms catalyzing As methylation in two anaerobic soil-derived microbiomes. The *arsM* gene can be expressed at similar, or slightly different levels in the absence or presence of As(III) in some organisms [28, 29], but expressed at significantly higher levels in the presence of As(III) in others [30–33]. Thus, we sought to identify *arsM* genes transcribed and ArsM proteins showing increased expression in the +As condition relative to the no-As control (Figure 3) but also irrespective of whether their expression was reported as increased relative to the no-As control (Figure S4).

**Figure 3.**
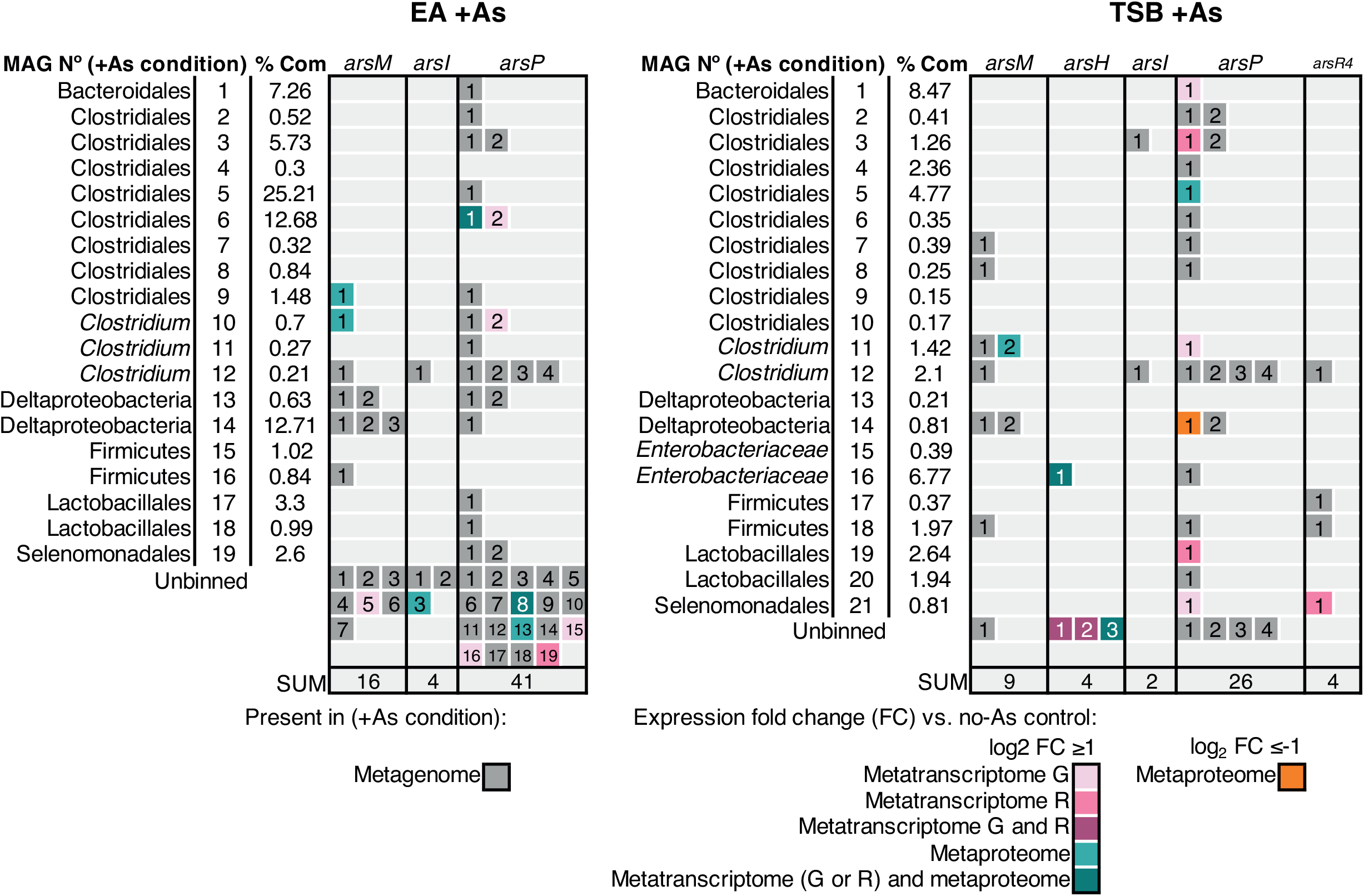
Distribution of *ars* genes involved in methylated arsenic metabolism encoded in MAGs from the +As condition and differentially expressed in metatranscriptomes/metaproteome relative to the no-As EA control. Each numbered box represents an *ars* gene. The number in each box corresponds to the “Numbering” column in Tables S2 and S12 where individual gene abundance and fold change values can be found. % Com: % Community as defined in caption from Table 1.

Sixteen phylogenetically distinct *arsM* genes were identified in the EA +As metagenome, but increased transcriptome reads or peptides (relative to the no-As control) were only detected for three genes (Figure 3) (see SI for details of calculation). The first is an *arsM* in Clostridiales EA MAG 9 classified by GhostKOALA as belonging to *Paeniclostridium sordellii* (EA MAG 9, *arsM-1*, psor type strain, in Table S2). The second was found in *Clostridium* EA MAG 10, also detected in the metaproteome, and the GhostKOALA taxonomic classification of the corresponding gene (EA MAG 10, *arsM-1* in Table S2) revealed that it was attributed to the unclassified species *Ruminococcaceae bacterium* CPB6 (Figure 3, Table S2), also referred as *Clostridium bacterium* CPB6 [34]. Finally, the third *arsM* was obtained from transcriptomic data but not clustered in any EA MAG (EA unbinned, *arsM-5* in Table S2) and likewise classified as pertaining to *Paeniclostridium sordellii*. Thus, in EA, all three *arsM* genes showing evidence of involvement in active As methylation pertain to fermenting microorganisms from the order Clostridiales.

Similarly, in the TSB +As metagenome, 9 distinct *arsM* genes were identified but none were detected in the metatranscriptome and only one exhibited increased expression in the metaproteome (Figure 3). It corresponds to an *arsM* gene from MAG 11 (TSB MAG11, *arsM-2* in Table S12). The expressed ArsM protein was assigned by GhostKOALA to a Clostridiales strain: *Clostridium botulinum* (*cby* type strain) (TSB MAG11, *arsM-2*) (Figure 3, Table S12). Finally, there was one *arsM* expressed in the TSB +As metaproteome but with no increased expression relative to the no-As control, it was classified as *Ruminococcaceae bacterium* CPB6 (TSB MAG 11, *arsM-1*) (Figure S4), the same organism identified in the EA culture (EA MAG 10, *arsM-1*). Thus, in TSB as in EA soil-derived microbiomes, As methylation appears to be catalyzed by various fermenting bacteria pertaining to the order Clostridiales such as members of the genera *Paeniclostridium* and *Clostridium*, and the family *Ruminococcaceae*. In addition to evidence for active arsenic methylation, there was evidence for active detoxification of methylated arsenic. Indeed, the metagenome included genes encoding proteins involved in the metabolism of methylated arsenic such as *arsH, arsI, arsP*, and *arsR4* (Figures 2 and 3). These genes encode proteins involved in the detoxification of methylated arsenic such as MMAs(III) and roxarsone: the oxidase ArsH, responsible for the oxidation of trivalent methylated As to the less toxic pentavalent form [35]; the demethylase ArsI that removes methyl groups from the As atom [36]; and the transmembrane transporter ArsP, thought to efflux methylated As [37]. The *arsR4* gene encodes an atypical MMAs(III)-responsive ArsR repressor, containing only two conserved cysteine residues [38]. The *Enterobacteriaceae* TSB MAG 16 exhibited activity of the oxygen-dependent ArsH protein [35] (Figure 3), a fact that is difficult to reconcile with the anoxic conditions. It is conceivable that this protein is capable of additional functions under anoxic conditions. An *arsR4*, shown to induce expression of *arsP* in the presence of MMAs(III) [38], had increased transcription along with an *arsP* encoded in the same contig in the Selenomonadales TSB MAG 21 (Figure 3, Table S12). Both genes transcripts were <5 TPM-RNA (Table S12) and thus, were not considered as transcribed in Figure S4. Finally, an ArsI protein, taxonomically related to class *Clostridia* ([*Eubacterium*] *rectale*), was expressed but it was encoded in an unbinned gene from the EA culture (Figure 3, Table S12). The identification of MAGs exhibiting a detoxification response to methylated As supports the hypothesis, raised above, of the role of monomethylated As as an arsenic antibiotic.

### Isolation of an arsenic-methylating anaerobic microorganism

Based on the analysis of the active metabolic activity from the EA MAG 9, expressing an As(III) methyltransferase (Figure S5), an appropriate selective medium was identified for its isolation. We utilized the fact that this MAG harbors and expresses the anaerobic assimilatory sulfite reductase encoded in the *asrABC* operon which is responsible for the NADH-dependent reduction of sulfite to sulfide [39–41] in sulfite-reducing Clostridia (SRC). It was the only member of Clostridiales expressing this capability in the EA microbiome (Figure S5). Thus, the isolation relied on growing the EA culture on agar medium selective for the SRC phenotype. In TSC agar, designed for the enumeration of *Clostridium perfringens* in food [42], the colonies from SRC are black, as the ammonium ferric citrate forms iron sulfide during sulfite reduction. Additionally, D-cycloserine acts a selective agent for the isolation of Clostridia strains [43] while inhibiting facultative anaerobes [42]. Finally, the bromocresol purple contained in the agar allows further differentiation between negative and positive sucrose fermenters, the latter changing the purple color of the agar to yellow. As none of the genes involved in sucrose transport or hydrolysis were binned in EA Clostridiales MAG 9 (Figure S5), only non-sucrose fermenting black colonies were considered. Those colonies were selected and using a colony PCR screen specifically targeting the *arsM* gene of EA MAG 9, we isolated a Clostridiales strain encoding the gene of the expressed ArsM in the EA MAG 9 (protein id k119_30669_28, Table S2) (Figure S6).

The isolate consists of non-sucrose-fermenting, rod-shaped and spore-forming bacteria forming convex and circular black colonies on TSC agar (Figures S7 and S8). The BLAST® (NCBI) search of the 16S rRNA sequence gives >99% identity to *Paraclostridium* strains (Table S13). On the basis of the 16S rRNA sequence, we assign the following name to the bacterium: “*Paraclostridium* species str. EML”. Strain EML was tested for As methylation under anaerobic conditions with 25 μM As(III). The growth of strain EML was hindered by As(III) (Figure 4-A) and starting from ~4 hours, the isolate transformed As(III) to monomethylated soluble As representing 48.3±1.5% of the soluble arsenic in the culture after 83 h (panels B and C from Figure 4). A fraction (14.7±0.6 μM) of the arsenic was found associated with biomass almost exclusively as inorganic arsenic (Figure 4-D).

**Figure 4.**
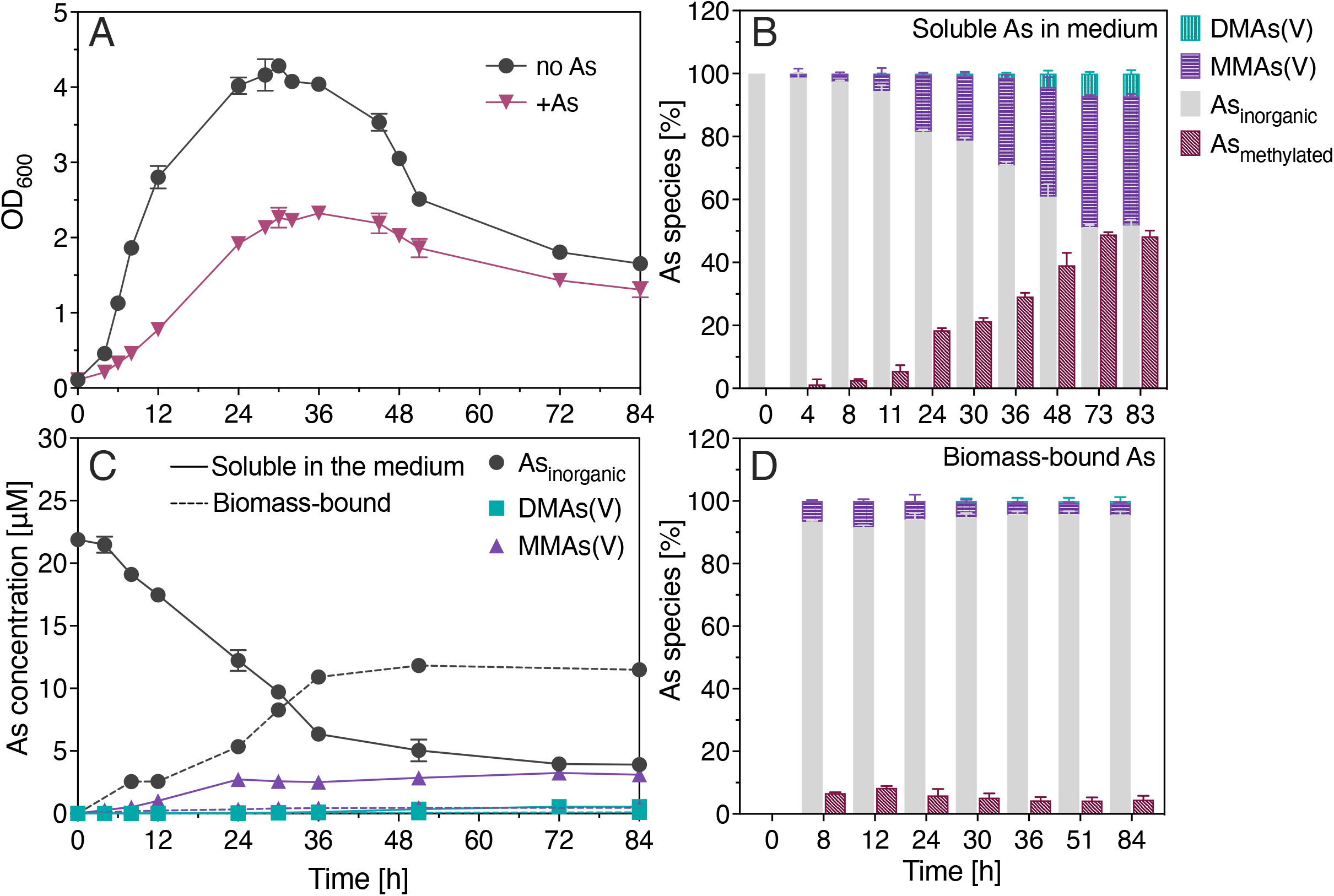
Isolate *“Paraclostridium* sp. EML”: (A) growth as OD_600_ with 25 μM As(III) and without, (B) percentage of soluble arsenic species in filtered medium containing 25 μM As(III), (C) concentration of arsenic species soluble in filtered medium containing 25 μM As(III) (solid lines) and biomass-bound (dashed lines) and (D) percentage of biomass-bound arsenic species. OD_600_ corresponds to optical density at 600 nm. Data points and bars represent the mean value and error bars, plus and minus one standard deviation. Individual values for each measurement and biological replicate are available in Tables S23 and S24.

## Discussion

Our results demonstrate the successful translation of multi-omic information to a specific strategy for targeted microbial isolation. While the metagenomes from the anaerobic soil-derived microbiomes identified the potential for As methylation in microorganisms from diverse taxa, the post-genomic approaches of community gene and protein expression clearly pointed to the active role of Clostridiales microorganisms in both cultures. This information paved the way for the identification of As-methylating microorganisms and the successful isolation of an anaerobic As methylator.

The EA and TSB soil-derived cultures offered the opportunity to study active As methylation from paddy-soil microbiomes in an environment that is less complex than soil but that remains environmentally relevant. In contrast to soil slurries, the lack of soil minerals facilitated the detection of soluble methylarsenicals and the facile extraction of DNA, RNA and proteins. The multi-omic approach made it possible to identify the putative microorganisms driving As methylation and their metabolism. Targeting a specific *arsM* gene rather than the synthesis of methylarsenicals greatly accelerated colony screening, as colony PCR could be employed instead of analytical detection of methylated As by HPLC-ICP-MS.

Had only the metagenomic approach been implemented, the data would have pointed to SRB MAGs as putative As methylators, as they harbored the most abundant *arsM* genes (Figure 6). Indeed, SRB have been proposed as drivers of As methylation in rice paddy soils based on the correlation in the abundance of the *arsM* and *dsrB* genes, and a decrease in As methylation by the addition chemical inhibitors of DSR [44, 45]. Yet, in the present findings, the SRB Deltaproteobacteria MAGs, although actively reducing sulfate (Figures S5 and S9), did not exhibit As-methylating activity as their *arsM* genes were neither transcribed nor translated (Figures 5). A recent study has shown that the abundance of *ars* genes in high-vs. low-As paddy soils was comparable whilst their transcriptomic activity was significantly impacted [46], bolstering the emerging view that to identify active As methylators in natural environments, the exclusive use of genomic data is insufficient.

**Figure 5.**
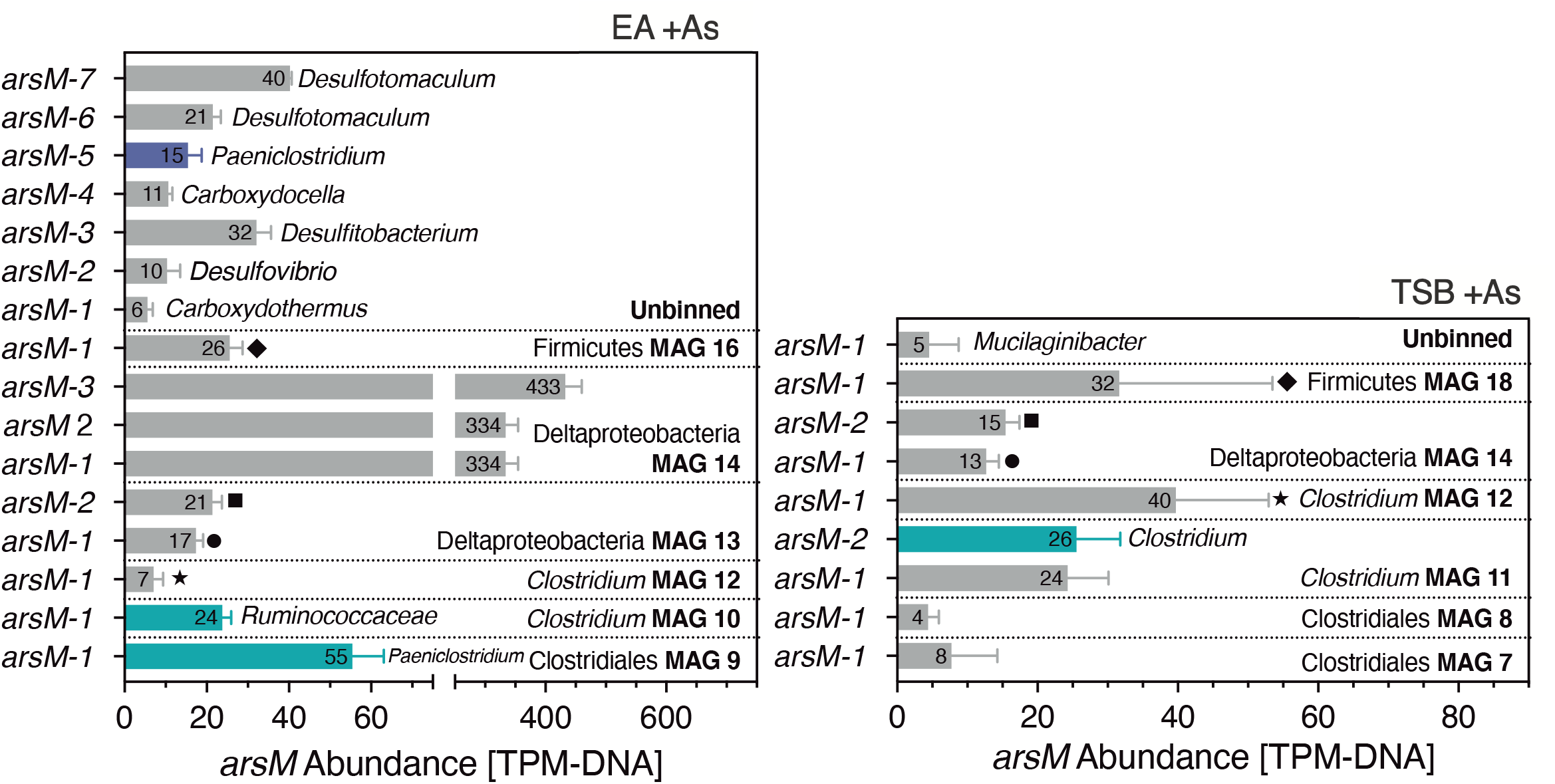
Gene abundance of *arsM* genes in MAGs from the +As condition cultures. Colored bars correspond to *arsM* genes with increased expression in the metaproteome (blue-green) or in the metatranscriptome G (purple) from +As condition relative to the no-As control in EA (left panel) and TSB (right panel) cultures. The taxonomic classification shown on the right side of the error bars for selected *arsM* genes corresponds to the individual gene classification assigned by GhostKOALA - “Genus” column in Tables S2 and S12. Columns with matching symbols on the right side of the error bars, correspond to matching *arsM* genes between the cultures. Individual gene abundance can be found in Tables S2 and S12. Numbers inside bar and bar length represent mean and error bars one standard deviation.

Previous work had identified another As-methylating Clostridiales strain, *Clostridium* sp. BXM [14], that performed fermentation and DSR. Amendment with organic matter [7, 47, 48], and an increase in dissolved organic carbon [49] have been shown to increase the As methylation efficiency, suggesting a correlation between the enrichment of fermenting communities, via increased availability of organic substrates, and As methylation. These observations along with the isolation of the present strain, point to a key role for fermenting Clostridiales microorganisms in As methylation.

It was previously proposed that the As-methylating activity of anaerobic microorganisms may be limited by efficient efflux of intracellular As(III) [13], or that it may function as a defensive response against nutrient competition [10]. However, up until now, the lack of anaerobic microbial isolates able to methylate As *in vitro* [13] precluded the investigation of this question. This study, capitalizing on the promise of omics-aided cultivation, paves the way for the elucidation of the controls on anaerobic As methylation. Further work is needed to elucidate why ArsM expression was restricted to members of Clostridiales fermenters and did not occur in other organisms harboring *arsM* genes.

## Supporting information

Supplemental Material&Methods and Results

## Acknowledgements

The work was funded by the Swiss National Science Foundation (SNSF) grant 310030_176146-1 and the SNSF NCCR Microbiomes (grant # 51NF40_180575). The work at ORNL was conducted under the Plant-Microbe Interface Science Focus Area, as supported by the U.S. Department of Energy, Office of Science, Office of Biological and Environmental Research, Genomic Science Program.

## Data availability

Metagenomic and metatranscriptomic raw sequencing reads are available at the National Centre for Biotechnology Information (NCBI) Sequence Read Archive (SRA) under BioProject PRJNA714492 (post publication). Data from the meta-omic analyses and source data from figures are available in Zenodo data repository (10.5281/zenodo.4605527, post publication) or by request from the authors.

## SI Figures Legends

**Figure S1. Arsenic methylation in the +As condition EA culture. A) Growth curves from experiments as OD_600_.** Red arrow indicates the samples used for metagenome and metaproteome analyses. Blue arrow indicates the samples used for metatranscriptomes G and R. B) Total soluble arsenic in medium. C) Percentage of arsenic species soluble in medium for metagenome and metaproteome experiments. D) Percentage of arsenic species soluble in medium for metatranscriptome G and metatranscriptome R experiments. OD_600_ corresponds to optical density at 600 nm. Points and bar heights represent mean and error bars plus, minus one standard deviation. Individual values for each measurement and biological replicate are available in Tables S25 and S26.

**Figure S2. Arsenic methylation in the +As condition TSB culture. A) Growth curves from experiments as OD_600_.** Red arrow indicates the samples used for metagenome and metaproteome analyses. Blue arrow indicates the samples used for metatranscriptomes G and R. B) Total soluble arsenic in medium. C) Percentage of arsenic species soluble in medium for metagenome and metaproteome experiments. D) Percentage of arsenic species soluble in medium for metatranscriptome G and metatranscriptome R experiments. OD_600_ corresponds to optical density at 600 nm. Points and bar heights represent mean and error bars plus, minus one standard deviation. Individual values for each measurement and biological replicate are available in Tables S25 and S26.

**Figure S3. Growth in no-As controls. Growth curves as OD_600_ from no-As controls in EA (left panel) and TSB (right panel) cultures.** Red arrow indicates the samples used for metagenome and metaproteome analyses. Blue arrow indicates the samples used for metatranscriptome G. OD_600_ corresponds to optical density at 600 nm. Points represent mean and error bars plus, minus one standard deviation. Individual values for biological replicate are available in Table S25.

**Figure S4. Distribution of *ars* genes involved in methylated arsenic metabolism encoded in MAGs from the +As condition and expressed in metatranscriptomes/metaproteome.** Each numbered box represents an *ars* gene. The number in each box corresponds to the “Numbering” column in Tables S2 and S12 where individual gene abundance and fold change values can be found. % Com: % Community as defined in caption from Table 1.

**Figure S5. Key enzymes from major metabolic pathways in MAGs from the +As condition EA culture.** Vertical bold lines correspond to the grouping of the MAGs with same lineage (Table 1-A). Pathway abbreviations: dissimilatory nitrate reduction to ammonia (DNRA), organic carbon metabolism (Org. C), propionic acid fermentation (propionic acid f.), and acetone-butanol-ethanol (ABE) fermentation. Refer to SI Table S16 for individual gene, transcript and protein abundance values and further enzymes.

**Figure S6. Colony PCR agarose gel from *Paraclostridium* sp. EML.** Left lane: PCR product from the amplification of *arsM* (protein id k119_30669_28, Table S2) in the PCR reaction using a *Paraclostridium* sp. EML colony, right lane: ladder corresponding to (from top to bottom) 1000, 750, 500, 300, 150 and 50 bp.

**Figure S7.** Light microscopy of *Paraclostridium* sp. EML cells, 48-h culture.

**Figure S8.** Growth of *Paraclostridium* sp. EML isolate in TSC agar.

**Figure S9. Key enzymes from major metabolic pathways in MAGs from the +As condition TSB culture.** Vertical bold lines correspond to the grouping of the MAGs with same lineage (Table 1-B). Pathway abbreviations: dissimilatory nitrate reduction to ammonia (DNRA), organic carbon metabolism (Org. C), propionic acid fermentation (propionic acid f.), and acetone-butanol-ethanol (ABE) fermentation. Refer to SI Table S17 for individual gene, transcript and protein abundance values and further enzymes.

**Figure S10. % Community of MAGs. % Community of MAGs in +As condition and no-As control from EA (top panel) and TSB (low panel) cultures.** Statistical differences between +As condition vs. no-As control were identified by unpaired Student t-test with p-value <0.05. Symbols: NA: no matching MAG in no-As control was found, one or more asterisks (*): significant difference and ns: no significant difference (p-value >0.05) (see Table S14 for P value symbol summary). Points represent individual values from three biological replicates. Bar heights represent mean and horizontal lines plus, minus one standard deviation.

**Figure S11. Volcano plots of metatranscriptomes.** Dots represent individual genes transcribed in metatranscriptomes R (upper panels) or G (lower panels) from +As condition EA (left panels) and TSB (right panels) cultures. Genes considered statistically differentially transcribed in the +As condition vs. no-As controls, based on the adjusted p value (q value), are represented as magenta (decreased transcription), green (increased transcription) and yellow (−1< log_2_ fold changes < 1) dots. Grey dots are genes with non-statistically significant changes in transcription. In bold numbers, the percentage of genes in magenta or green. Individual fold-change values are available in Tables S19 and S20.

**Figure S12. Volcano plots of metaproteomes.** Dots represent individual genes expressed in metaproteomes from +As condition EA (left panel) and TSB (right panel) cultures. Genes considered statistically differentially expressed in the +As condition vs. respective no-As controls, based on the adjusted p value (q value), are represented as magenta (decreased expression), green (increased expression) and yellow (−1< log_2_ fold changes < 1) dots. Grey dots are genes with non-statistically significant changes in expression. In bold numbers, the percentage of genes in magenta or green. Individual fold-change values are available in Tables S19 and S20.

**Figure S13. Distribution of *ars* genes encoded in MAGs from the +As condition EA culture and differentially expressed in metatranscriptomes/metaproteome relative to the no-As EA control.** Each numbered box represents an *ars* gene. The number in each box corresponds to the “Numbering” column in Table S2 where individual gene abundance and fold change values can be found.

**Figure S14. Distribution of *ars* genes encoded in MAGs from the +As condition TSB culture and differentially expressed in metatranscriptomes/metaproteome relative to the no-As TSB control.** Each numbered box represents an *ars* gene. The number in each box corresponds to the “Numbering” column in Table S12 where individual gene abundance and fold change values can be found.

**Figure S15. Edwards-Venn diagrams of *ars* genes with increased expression in +As condition EA and TSB relative to no-As control cultures.** Number of *ars* genes encoded in metagenomes, with increased expression in metatranscriptomes R and G or/and metaproteomes from +As condition EA culture (left panel) and +As condition TSB (right panel) cultures.

## References

1. Zhu Y-G, Yoshinaga M, Zhao F-J, Rosen BP. Earth Abides Arsenic Biotransformations. Annu Rev Earth Planet Sci 2014; 42: 443–467.

2. Lomax C, Liu WJ, Wu L, Xue K, Xiong J, Zhou J, et al. Methylated arsenic species in plants originate from soil microorganisms. New Phytol 2012; 193: 665–672.

3. Zhao FJ, Zhu YG, Meharg AA. Methylated arsenic species in rice: Geographical variation, origin, and uptake mechanisms. Environ Sci Technol 2013; 47: 3957–3966.

4. Zheng MZ, Li G, Sun GX, Shim H, Cai C. Differential toxicity and accumulation of inorganic and methylated arsenic in rice. Plant Soil 2013; 365: 227–238.

5. Abedin MJ, Feldmann J, Meharg AA. Uptake kinetics of arsenic species in rice plants. Plant Physiol 2002; 128: 1120–1128.

6. Dunivin TK, Yeh SY, Shade A. A global survey of arsenic-related genes in soil microbiomes. BMC Biol 2019; 17.

7. Jia Y, Huang H, Zhong M, Wang F-H, Zhang L-M, Zhu Y-G. Microbial Arsenic Methylation in Soil and Rice Rhizosphere. Environ Sci Technol 2013; 47: 3141–3148.

8. Xiao KQ, Li LG, Ma LP, Zhang SY, Bao P, Zhang T, et al. Metagenomic analysis revealed highly diverse microbial arsenic metabolism genes in paddy soils with low-arsenic contents. Environ Pollut 2016; 211: 1–8.

9. Zhang SY, Su JQ, Sun GX, Yang Y, Zhao Y, Ding J, et al. Land scale biogeography of arsenic biotransformation genes in estuarine wetland. Environ Microbiol 2017; 19: 2468–2482.

10. Chen J, Rosen BP. The Arsenic Methylation Cycle: How Microbial Communities Adapted Methylarsenicals for Use as Weapons in the Continuing War for Dominance. Front Environ Sci. 2020. Frontiers Media S.A., 8: 43

11. Yang Y, Wu S, Lilley RM, Zhang R. The diversity of membrane transporters encoded in bacterial arsenic-resistance operons. PeerJ 2015; 3: e943.

12. Yang P, Ke C, Zhao C, Kuang Q, Liu B, Xue X, et al. ArsM-mediated arsenite volatilization is limited by efflux catalyzed by As efflux transporters. Chemosphere 2020; 239: 124822.

13. Viacava K, Meibom KL, Ortega D, Dyer S, Gelb A, Falquet L, et al. Variability in Arsenic Methylation Efficiency across Aerobic and Anaerobic Microorganisms. Environ Sci Technol 2020; 54: 14343–14351.

14. Wang PP, Bao P, Sun GX. Identification And Catalytic Residues Of The Arsenite Methyltransferase From a Sulfate-Reducing Bacterium, Clostridium sp. BXM. FEMS Microbiol Lett 2015; 362: 1–8.

15. Reid MC, Maillard J, Bagnoud A, Falquet L, Le Vo P, Bernier-Latmani R. Arsenic Methylation Dynamics in a Rice Paddy Soil Anaerobic Enrichment Culture. Environ Sci Technol 2017; 51: 10546–10554.

16. Bright DA, Brock S, Reimer KJ, Cullen WR, Hewitt GM, Jafaar J. Methylation of arsenic by anaerobic microbial consortia isolated from lake sediment. Appl Organomet Chem 1994; 8: 415–422.

17. DeLong EF. Archaea in coastal marine environments. Proc Natl Acad Sci U S A 1992; 89: 5685–5689.

18. Edwards U, Rogall T, Blöcker H, Emde M, Böttger EC. Isolation and direct complete nucleotide determination of entire genes. Characterization of a gene coding for 16S ribosomal RNA. Nucleic Acids Res 1989; 17: 7843–7853.

19. Wang PP, Sun GX, Zhu YG. Identification And Characterization Of Arsenite Methyltransferase From An Archaeon, Methanosarcina Acetivorans C2A. Environ Sci Technol 2014; 48: 12706–12713.

20. Parks DH, Imelfort M, Skennerton CT, Hugenholtz P, Tyson GW. CheckM: Assessing the quality of microbial genomes recovered from isolates, single cells, and metagenomes. Genome Res 2015; 25: 1043–1055.

21. Lin Y-F, Walmsley AR, Rosen BP. An arsenic metallochaperone for an arsenic detoxification pump. Proc Natl Acad Sci U S A 2006; 103: 15617–22.

22. Shi W, Wu J, Rosen BP. Identification of a putative metal binding site in a new family of metalloregulatory proteins. J Biol Chem 1994; 269: 19826–19829.

23. Murphy JN, Saltikov CW. The ArsR repressor mediates arsenite-dependent regulation of arsenate respiration and detoxification operons of Shewanella sp. strain ANA-3. J Bacteriol 2009; 191: 6722–6731.

24. Santha S, Pandaranayaka EPJ, Rosen BP, Thiyagarajan S. Purification, crystallization and preliminary X-ray diffraction studies of the arsenic repressor ArsR from Corynebacterium glutamicum. Acta Crystallogr Sect F Struct Biol Cryst Commun 2011; 67: 1616–1618.

25. Fekih I Ben, Zhang C, Li YP, Zhao Y, Alwathnani HA, Saquib Q, et al. Distribution of arsenic resistance genes in prokaryotes. Front Microbiol 2018; 9: 2473.

26. Cai L, Yu K, Yang Y, Chen BW, Li XD, Zhang T. Metagenomic exploration reveals high levels of microbial arsenic metabolism genes in activated sludge and coastal sediments. Appl Microbiol Biotechnol 2013; 97: 9579–9588.

27. Zhang SY, Zhao FJ, Sun GX, Su JQ, Yang XR, Li H, et al. Diversity and abundance of arsenic biotransformation genes in paddy soils from Southern China. Environ Sci Technol 2015; 49: 4138–4146.

28. Huang K, Xu Y, Packianathan C, Gao F, Chen C, Zhang J, et al. Arsenic methylation by a novel ArsM As(III) S-adenosylmethionine methyltransferase that requires only two conserved cysteine residues. Mol Microbiol 2018; 107: 265–276.

29. Zhang J, Cao T, Tang Z, Shen Q, Rosen BP, Zhao FJ. Arsenic methylation and volatilization by arsenite s-adenosylmethionine methyltransferase in Pseudomonas alcaligenes NBRC14159. Appl Environ Microbiol 2015;81: 2852–2860.

30. Huang K, Chen C, Zhang J, Tang Z, Shen Q, Rosen BP, et al. Efficient Arsenic Methylation and Volatilization Mediated by a Novel Bacterium from an Arsenic-Contaminated Paddy Soil. Environ Sci Technol 2016; 50: 6389–6396.

31. Yin XX, Chen J, Qin J, Sun GX, Rosen BP, Zhu YG. Biotransformation and volatilization of arsenic by three photosynthetic cyanobacteria. Plant Physiol 2011; 156: 1631–1638.

32. Zhao C, Zhang Y, Chan Z, Chen S, Yang S. Insights into arsenic multi-operons expression and resistance mechanisms in Rhodopseudomonas palustris CGA009. Front Microbiol 2015; 6: 986.

33. Wang G, Kennedy SP, Fasiludeen S, Rensing C, DasSarma S. Arsenic Resistance in Halobacterium sp. Strain NRC-1 Examined by Using an Improved Gene Knockout System. J Bacteriol 2004; 186: 3187–3194.

34. Subhraveti P, Ong Q, Keseler I, Kothari A, Caspi R, Karp PD. Summary of Clostridium sp. CPB-6 CPB6, version 23.5. https://biocyc.org/organism-summary?object=GCF_002119605. Accessed 30 Mar 2020.

35. Chen J, Bhattacharjee H, Rosen BP. ArsH is an organoarsenical oxidase that confers resistance to trivalent forms of the herbicide monosodium methylarsenate and the poultry growth promoter roxarsone. Mol Microbiol 2015; 96: 1042–1052.

36. Yoshinaga M, Rosen BP. A C-As lyase for degradation of environmental organoarsenical herbicides and animal husbandry growth promoters. Proc Natl Acad Sci U S A 2014; 111: 7701–6.

37. Chen J, Madegowda M, Bhattacharjee H, Rosen BP. ArsP: A methylarsenite efflux permease. Mol Microbiol 2015; 98: 625–635.

38. Chen J, Nadar VS, Rosen BP. A novel MAs(III)-selective ArsR transcriptional repressor. Mol Microbiol 2017; 106: 469–478.

39. Doyle CJ, O’Toole PW, Cotter PD. Genomic characterization of sulphite reducing bacteria isolated from the dairy production chain. Front Microbiol 2018; 9: 1507.

40. Czyzewski BK, Wang DN. Identification and characterization of a bacterial hydrosulphide ion channel. Nature 2012; 483: 494–497.

41. Huang CJ, Barrett EL. Sequence analysis and expression of the Salmonella typhimurium asr operon encoding production of hydrogen sulfide from sulfite. J Bacteriol 1991; 173: 1544–1553.

42. Harmon SM, Kautter DA, Peeler JT. Improved medium for enumeration of Clostridium perfringens. Appl Microbiol 1971; 22: 688–692.

43. George WL, Sutter VL, Finegold SM. Toxigenicity and antimicrobial susceptibility of Clostridium difficile, a cause of antimicrobial agent-associated colitis. Curr Microbiol 1978; 1: 55–58.

44. Chen C, Li L, Huang K, Zhang J, Xie WY, Lu Y, et al. Sulfate-reducing bacteria and methanogens are involved in arsenic methylation and demethylation in paddy soils. ISME J 2019; 13: 2523–2535.

45. Wang M, Tang Z, Chen XP, Wang X, Zhou WX, Tang Z, et al. Water management impacts the soil microbial communities and total arsenic and methylated arsenicals in rice grains. Environ Pollut 2019; 247: 736–744.

46. Zhang S-Y, Xiao X, Chen S-C, Zhu Y-G, Sun G-X, Konstantinidis KT. High arsenic levels increase activity rather than diversity or abundance of arsenic metabolism genes in paddy soils. Appl Environ Microbiol 2021.

47. Mestrot A, Feldmann J, Krupp EM, Hossain MS, Roman-Ross G, Meharg AA. Field fluxes and speciation of arsines emanating from soils. Environ Sci Technol 2011; 45: 1798–1804.

48. Huang H, Jia Y, Sun G-X, Zhu Y-G. Arsenic speciation and volatilization from flooded paddy soils amended with different organic matters. Environ Sci Technol 2012; 46: 2163–8.

49. Zhao F-J, Harris E, Yan J, Ma J, Wu L, Liu W, et al. Arsenic Methylation in Soils and Its Relationship with Microbial arsM Abundance and Diversity, and As Speciation in Rice. Environ Sci Technol 2013; 7147–7154.

