## Supplemental Figures for "Meta-omics-aided isolation of an elusive anaerobic arsenic-methylating soil bacterium"

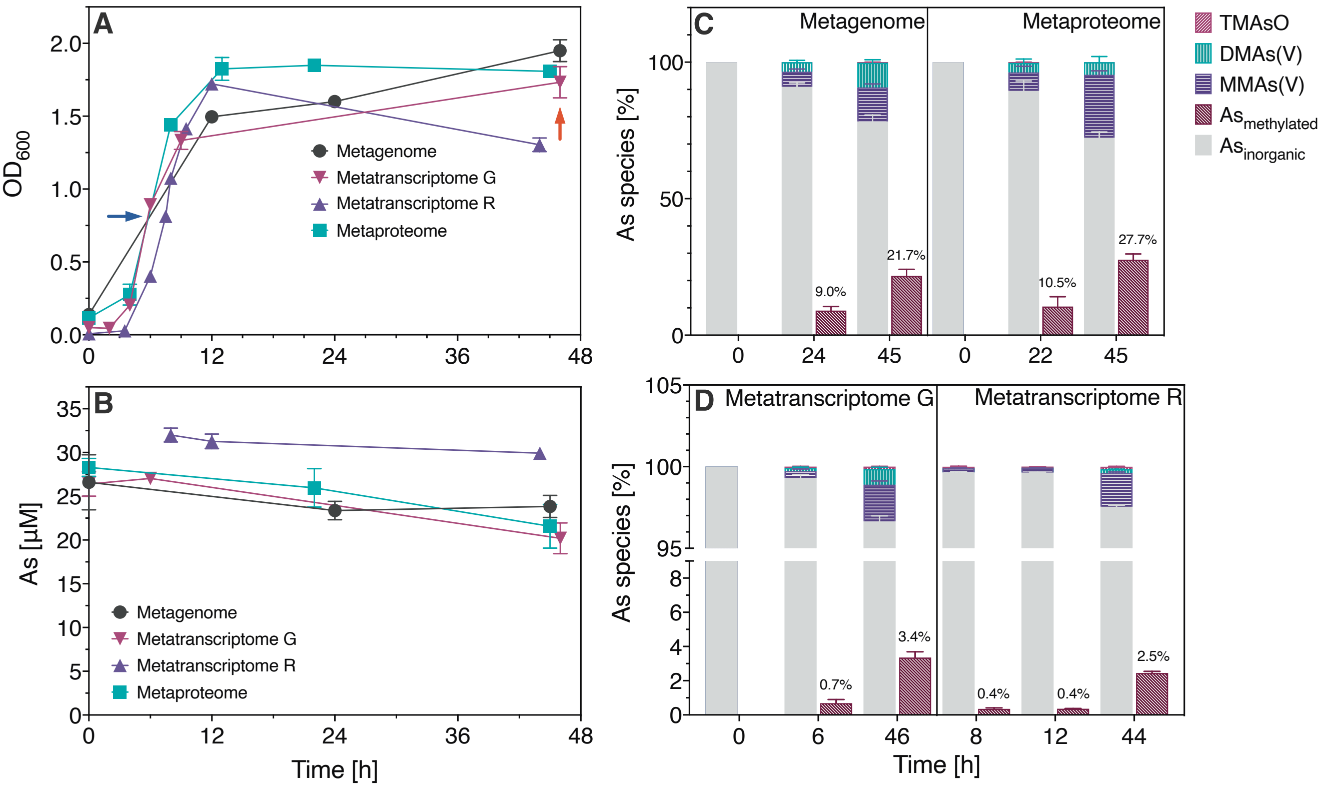


**Figure S1. Arsenic methylation in the +As condition EA culture.** A) Growth curves from experiments as $OD_{600}$. Red arrow indicates the samples used for metagenome and metaproteome analyses. Blue arrow indicates the samples used for metatranscriptomes G and R. B) Total soluble arsenic in medium. C) Percentage of arsenic species soluble in medium for metagenome and metaproteome experiments. D) Percentage of arsenic species soluble in medium for metatranscriptome G and metatranscriptome R experiments. $OD_{600}$ corresponds to optical density at 600 nm. Points and bar heights represent mean and error bars plus, minus one standard deviation. Individual values for each measurement and biological replicate are available in Tables S25 and S26.

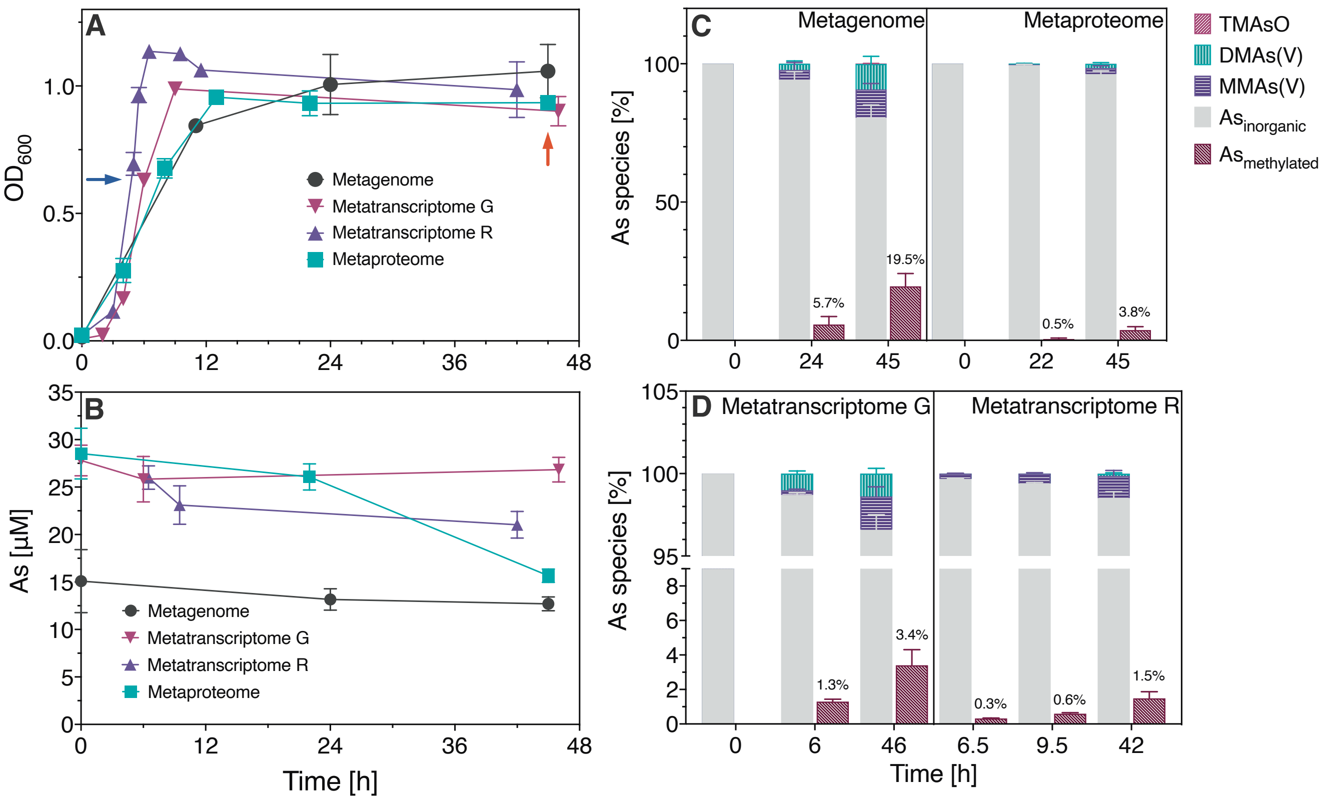


**Figure S2. Arsenic methylation in the +As condition TSB culture.** A) Growth curves from experiments as $OD_{600}$. Red arrow indicates the samples used for metagenome and metaproteome analyses. Blue arrow indicates the samples used for metatranscriptomes G and R. B) Total soluble arsenic in medium. C) Percentage of arsenic species soluble in medium for metagenome and metaproteome experiments. D) Percentage of arsenic species soluble in medium for metatranscriptome G and metatranscriptome R experiments. $OD_{600}$ corresponds to optical density at 600 nm. Points and bar heights represent mean and error bars plus, minus one standard deviation. Individual values for each measurement and biological replicate are available in Tables S25 and S26.

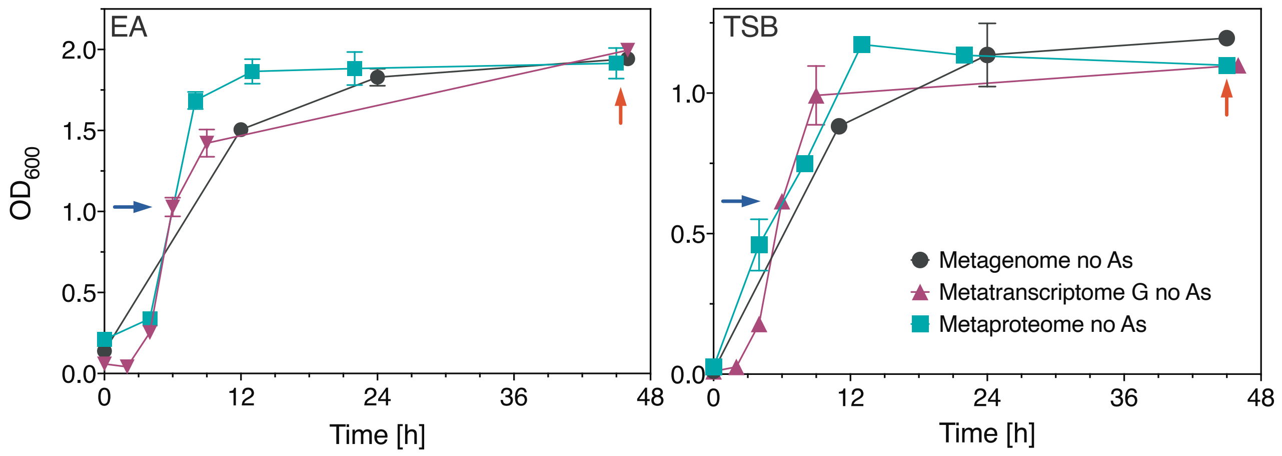


**Figure S3. Growth in no-As controls.** Growth curves as $OD_{600}$ from no-As controls in EA (left panel) and TSB (right panel) cultures. Red arrow indicates the samples used for metagenome and metaproteome analyses. Blue arrow indicates the samples used for metatranscriptome G. Points represent mean and error bars plus, minus one standard deviation. Individual values for biological replicate are available in Table S25.

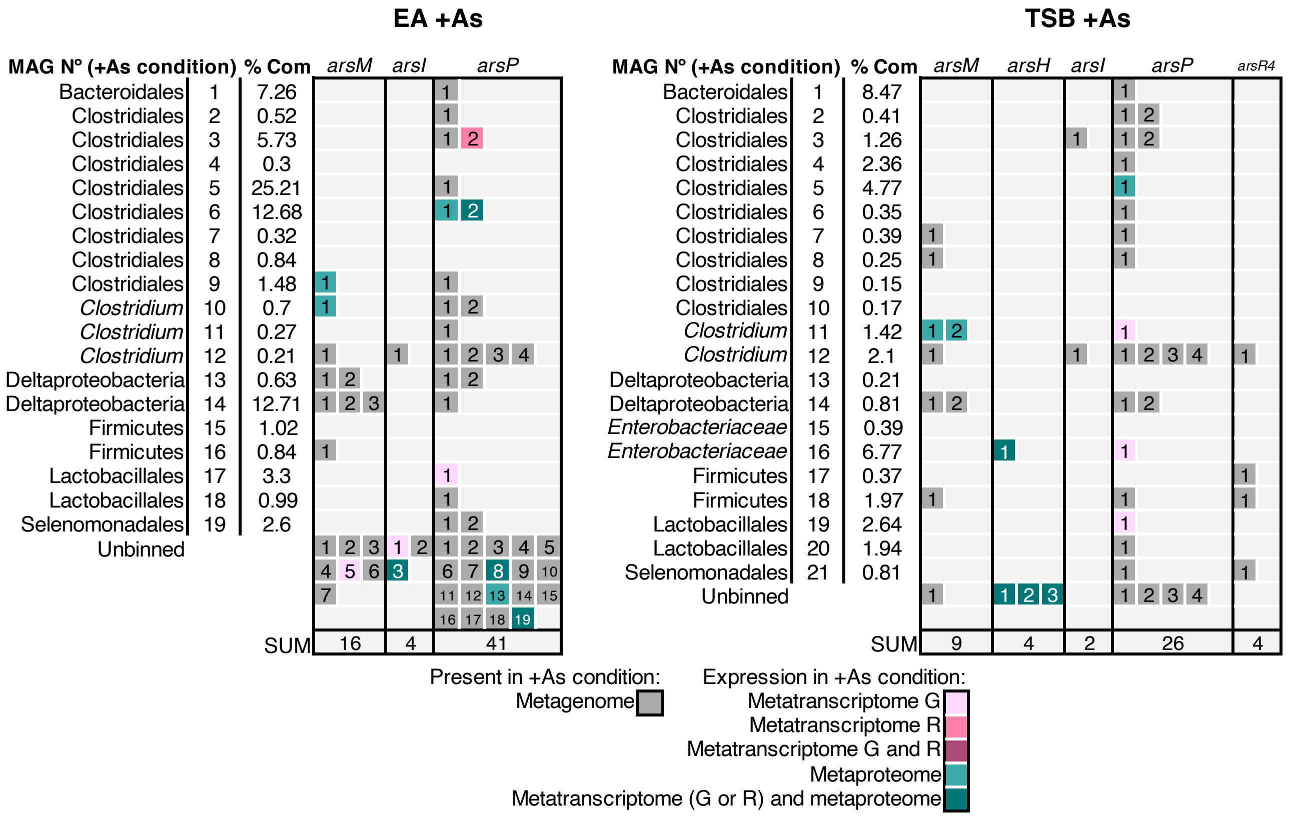


**Figure S4. Distribution of *ars* genes involved in methylated arsenic metabolism encoded in MAGs from the +As condition and expressed in metatranscriptomes/metaproteome.** Each numbered box represents an *ars* gene. The number in each box corresponds to the “Numbering” column in Tables S2 and S12 where individual gene abundance and fold change values can be found. % Com: % Community as defined in caption from Tables 1 and 2.

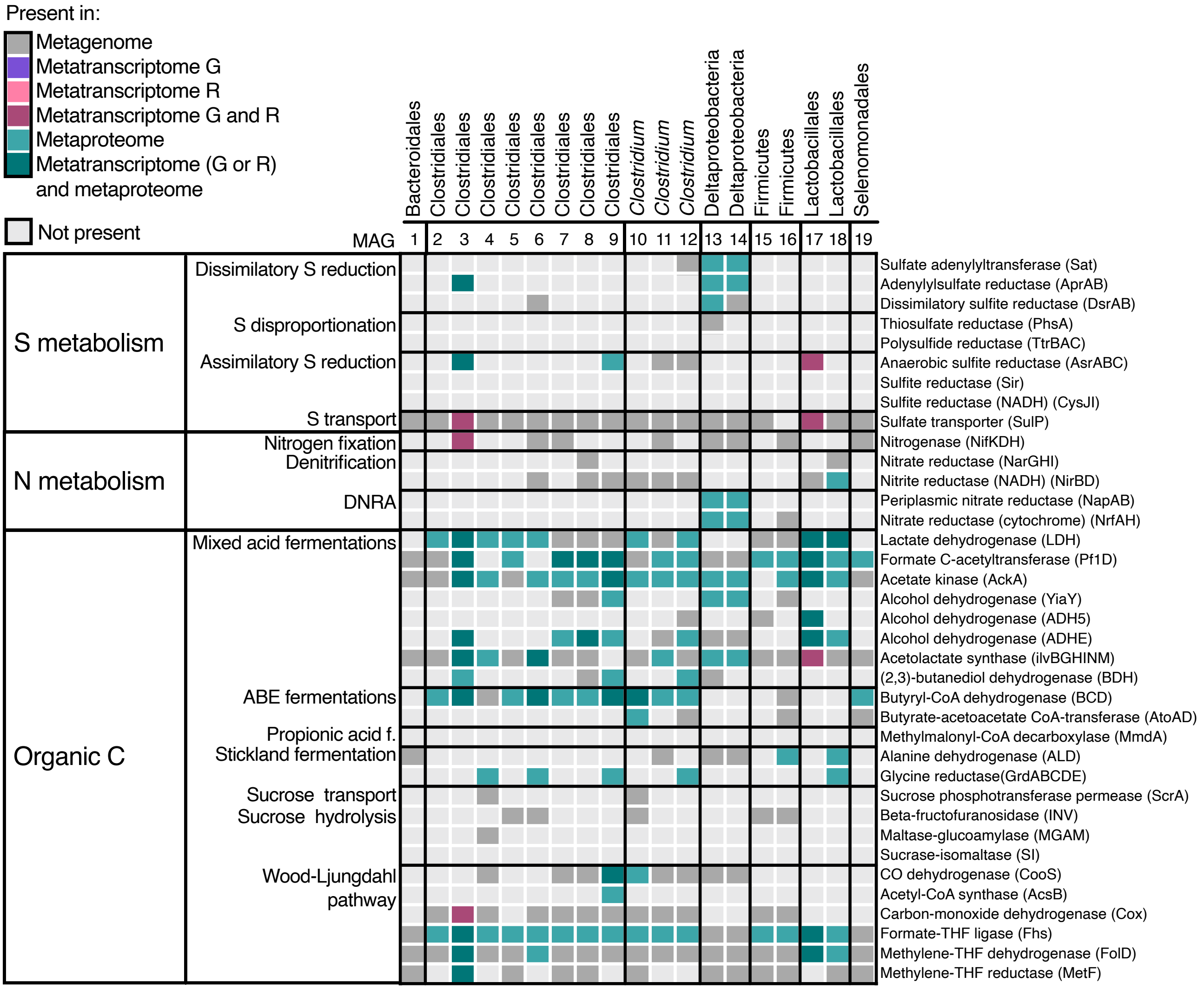


**Figure S5. Key enzymes from major metabolic pathways in MAGs from the +As condition EA culture.** Vertical bold lines correspond to the grouping of the MAGs with same lineage (Table 1). Pathway abbreviations: dissimilatory nitrate reduction to ammonia (DNRA), organic carbon metabolism (Org. C), propionic acid fermentation (propionic acid f.), and acetone-butanol-ethanol (ABE) fermentation. Refer to SI Table S16 for individual gene, transcript and protein abundance values and further enzymes.

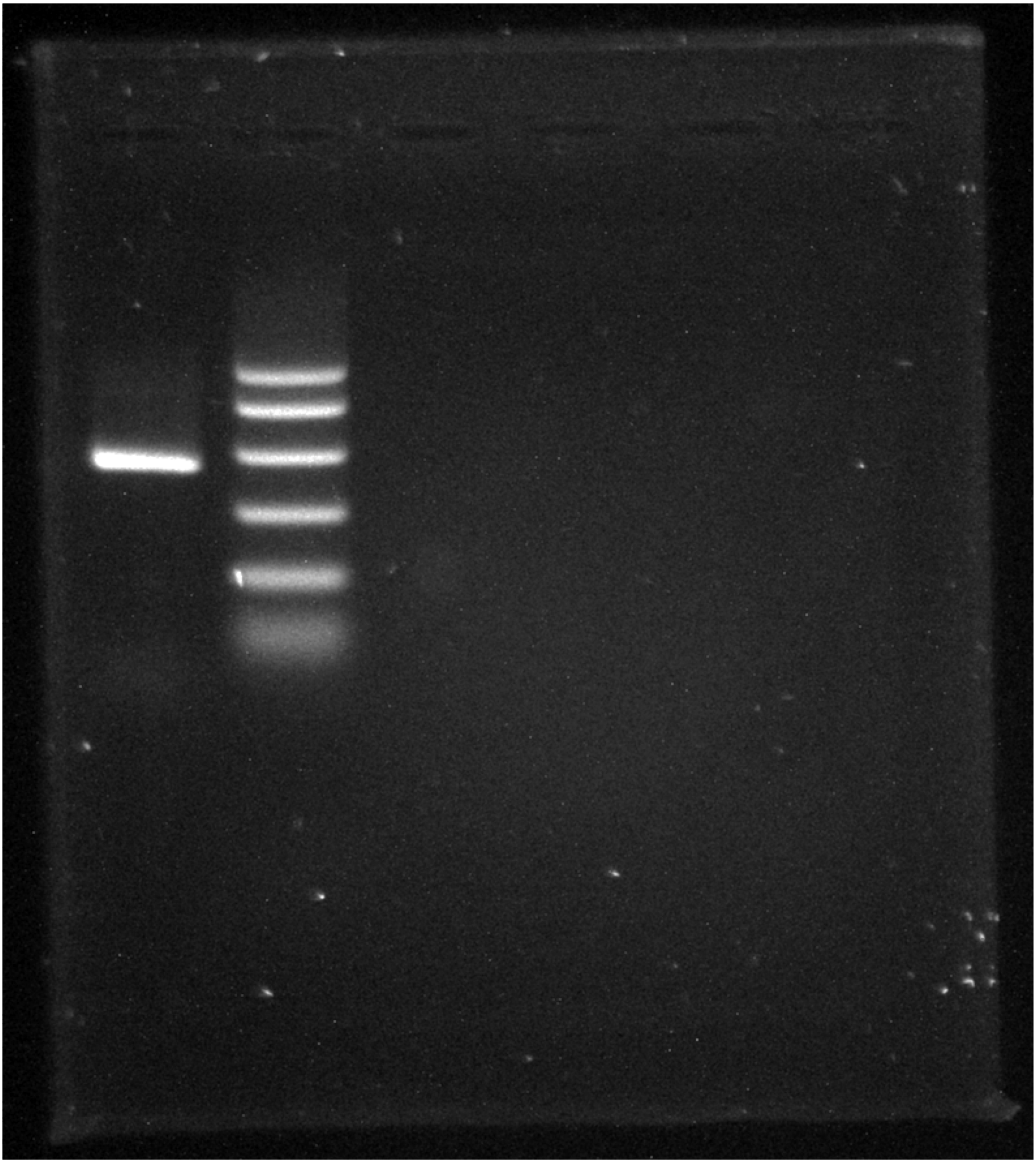

**Figure S6. Colony PCR agarose gel from *Paraclostridium* sp. EML.** Left lane: PCR product from the amplification of *arsM* (protein id k119_30669_28, Table S2) in the PCR reaction using a *Paraclostridium* sp. EML colony, right lane: ladder corresponding to (from top to bottom) 1000, 750, 500, 300, 150 and 50 bp.

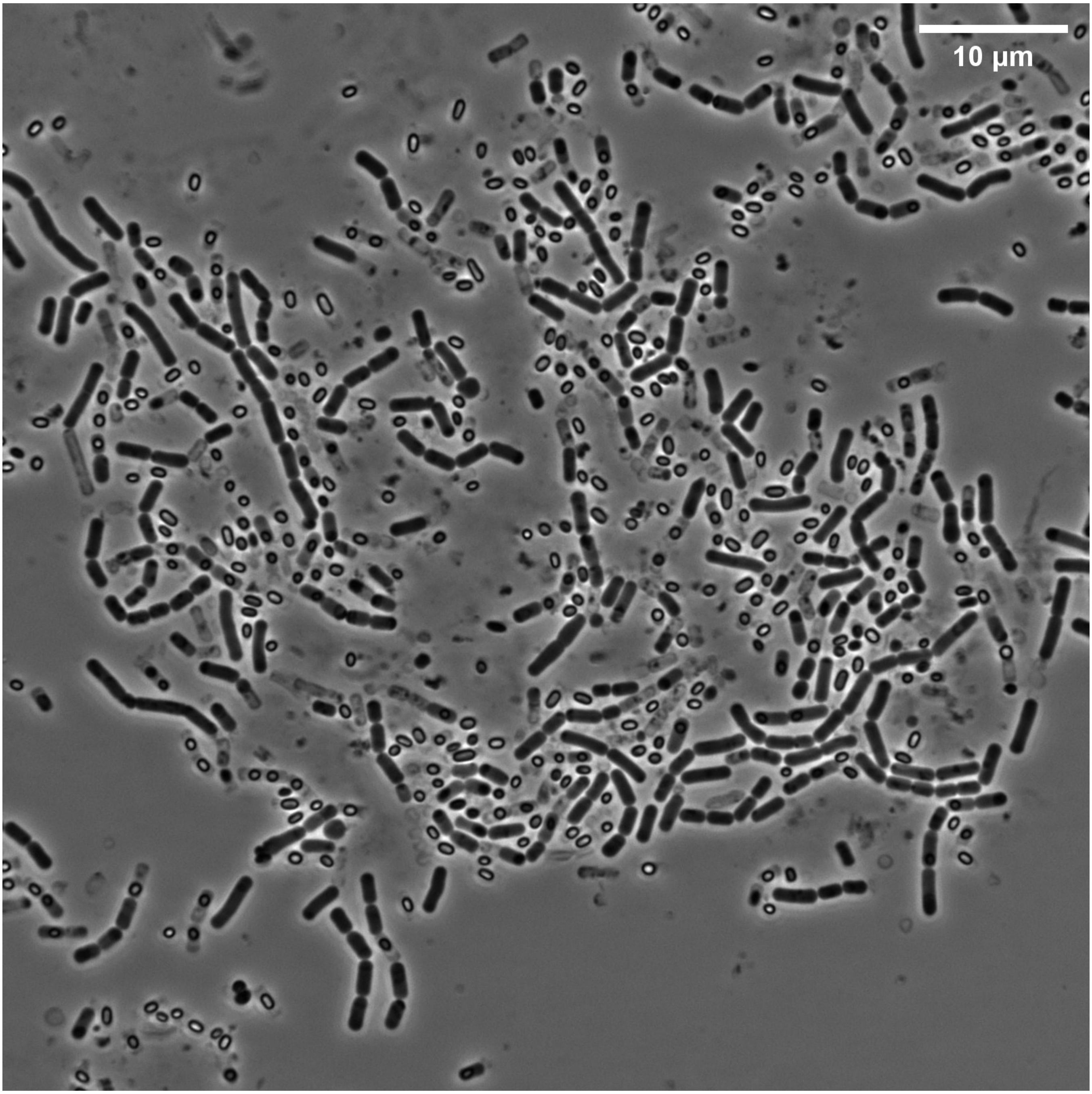


**Figure S7. Light microscopy of *Paraclostridium* sp. EML cells, 48-h culture.**

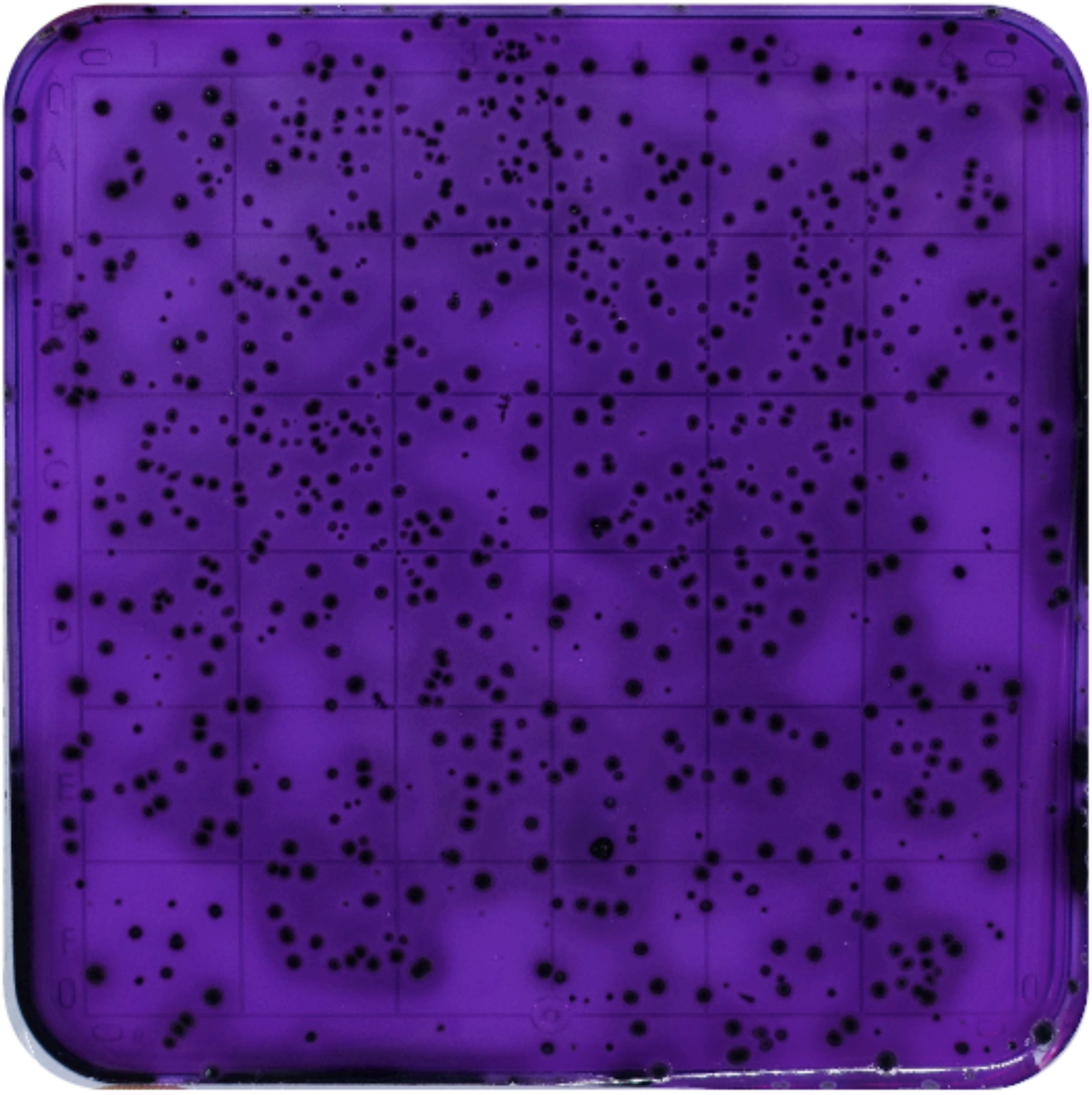

**Figure S8. Growth of *Paraclostridium* sp. EML isolate in TSC agar.**

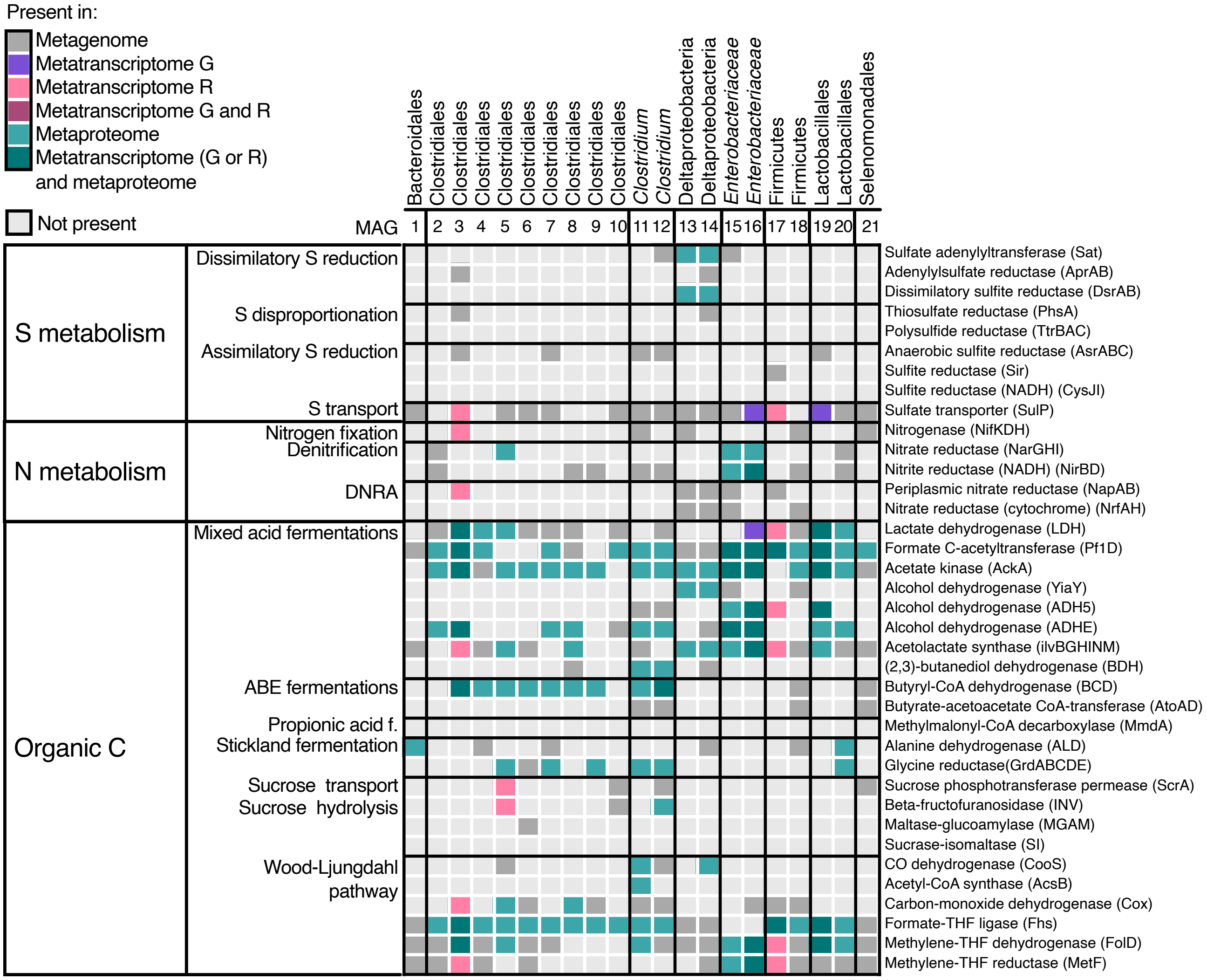


**Figure S9. Key enzymes from major metabolic pathways in MAGs from the +As condition TSB culture.** Vertical bold lines correspond to the grouping of the MAGs with same lineage (Table 2). Pathway abbreviations: dissimilatory nitrate reduction to ammonia (DNRA), organic carbon metabolism (Org. C), propionic acid fermentation (propionic acid f.), and acetone-butanol-ethanol (ABE) fermentation. Refer to SI Table S17 for individual gene, transcript and protein abundance values and further enzymes.

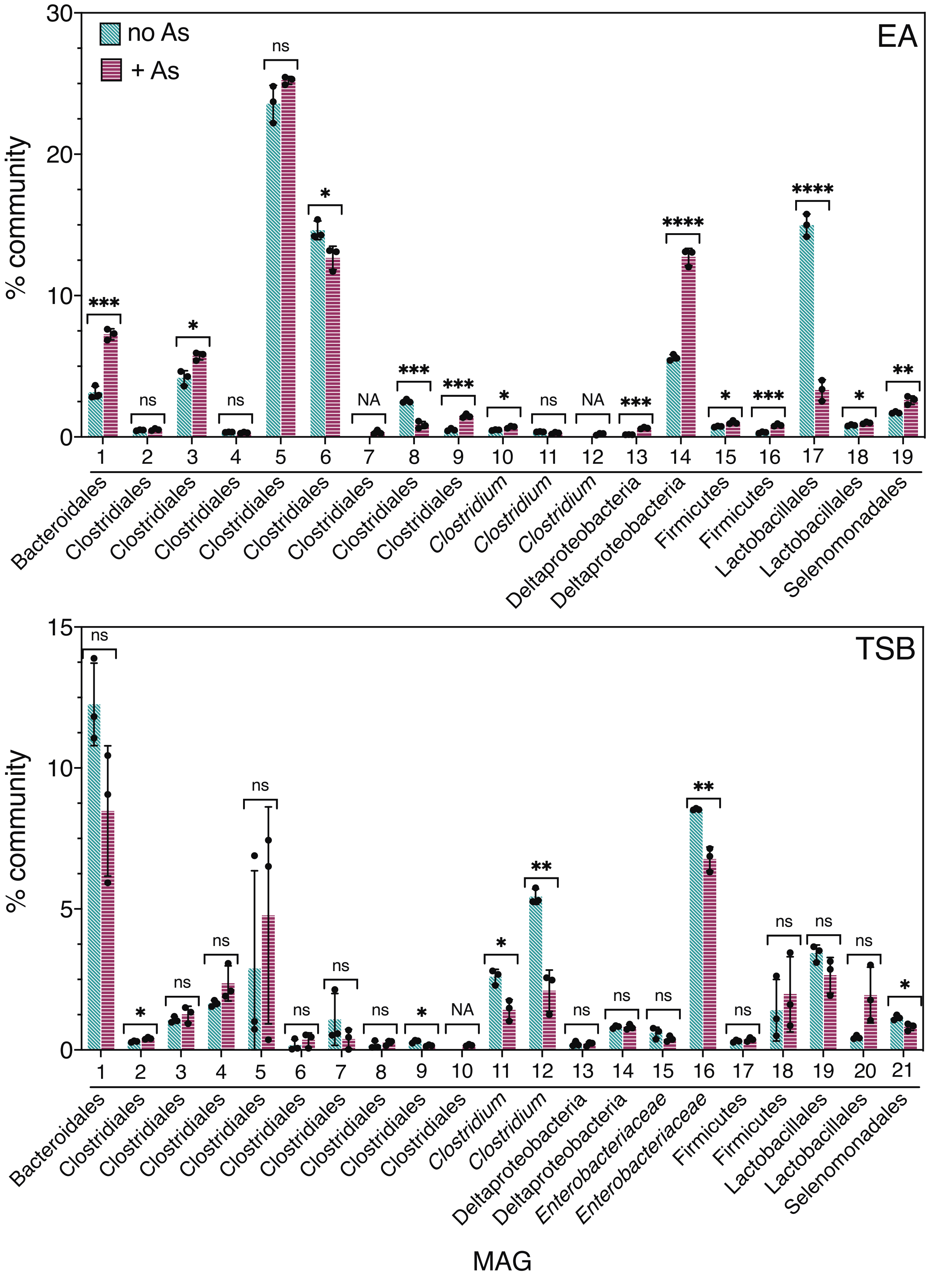


**Figure S10. % Community of MAGs.** % Community of MAGs in +As condition and no-As control from EA (top panel) and TSB (low panel) cultures. Statistical differences between +As condition vs. no-As control were identified by unpaired Student t-test with p-value <0.05. Symbols: NA: no matching MAG in no-As control was found, one or more asterisks (*): significant difference and ns: no significant difference (p-value >0.05) (see Table S14 for P value symbol summary) . Points represent individual values from three biological replicates. Bar heights represent mean and horizontal lines plus, minus one standard deviation.

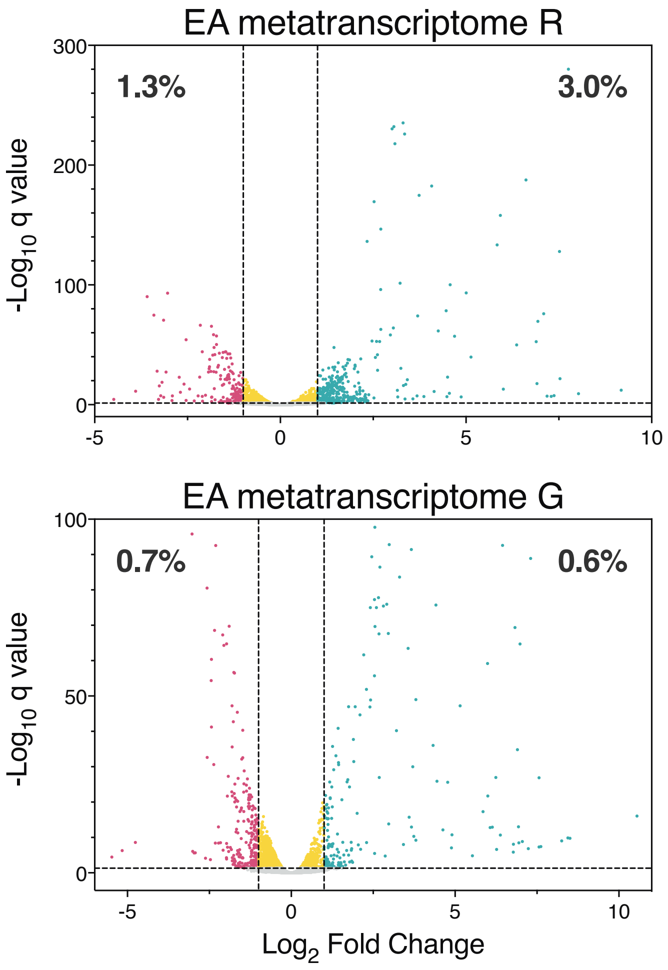


**Figure S11. Volcano plots of metatranscriptomes.** Dots represent individual genes transcribed in metatranscriptomes R (upper panels) or G (lower panels) from +As condition EA (left panels) and TSB (right panels) cultures. Genes considered statistically differentially transcribed in the +As condition vs. no-As controls, based on the adjusted p value (q value), are represented as magenta (decreased transcription), green (increased transcription) and yellow ($-1 < \log_2$ fold changes $< 1$) dots. Grey dots are genes with non-statistically significant changes in transcription. In bold numbers, the percentage of genes in magenta or green. Individual fold-change values are available in Tables S19 and S20.

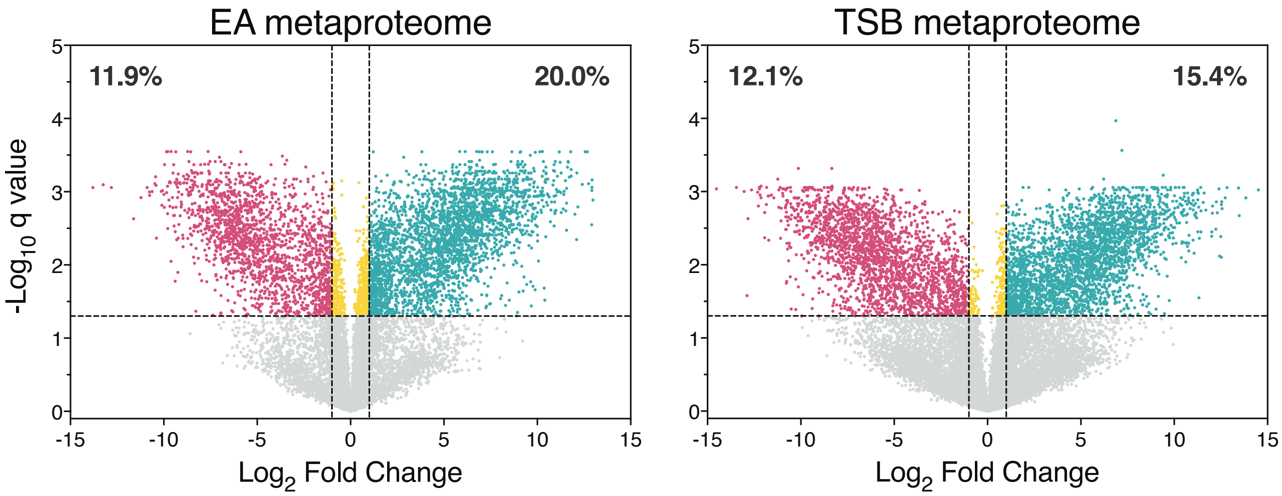


**Figure S12. Volcano plots of metaproteomes.** Dots represent individual genes expressed in metaproteomes from +As condition EA (left panel) and TSB (right panel) cultures. Genes considered statistically differentially expressed in the +As condition vs. respective no-As controls, based on the adjusted p value (q value), are represented as magenta (decreased expression), green (increased expression) and yellow ($-1 < \log_2$ fold changes $< 1$) dots. Grey dots are genes with non-statistically significant changes in expression. In bold numbers, the percentage of genes in magenta or green. Individual fold-change values are available in Tables S19 and S20.

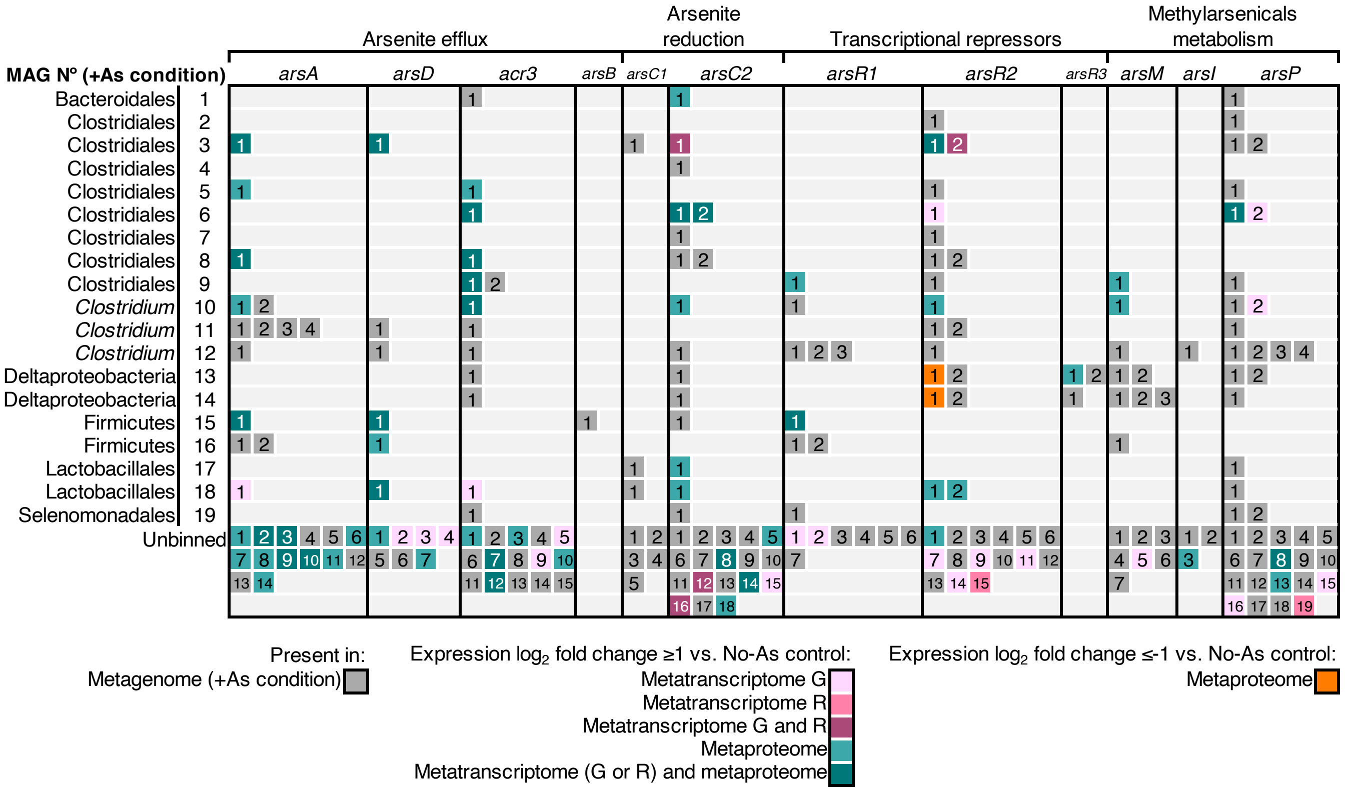


**Figure S13. Distribution of *ars* genes encoded in MAGs from the +As condition EA culture and differentially expressed in metatranscriptomes/metaproteome relative to the no-As EA control.** Each numbered box represents an *ars* gene. The number in each box corresponds to the "Numbering" column in Table S2 where individual gene abundance and fold change values can be found.

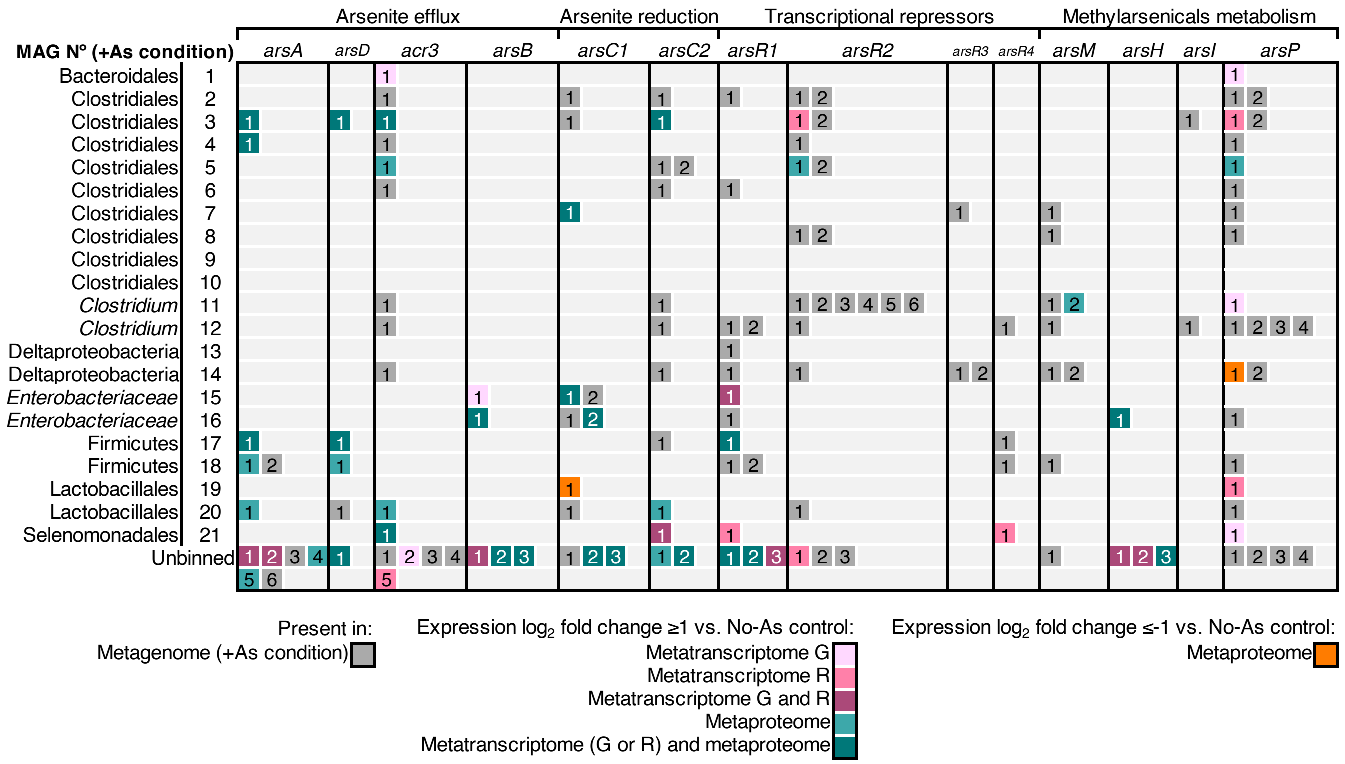


**Figure S14. Distribution of *ars* genes encoded in MAGs from the +As condition TSB culture and differentially expressed in metatranscriptomes/metaproteome relative to the no-As TSB control.** Each numbered box represents an *ars* gene. The number in each box corresponds to the "Numbering" column in Table S12 where individual gene abundance and fold change values can be found.

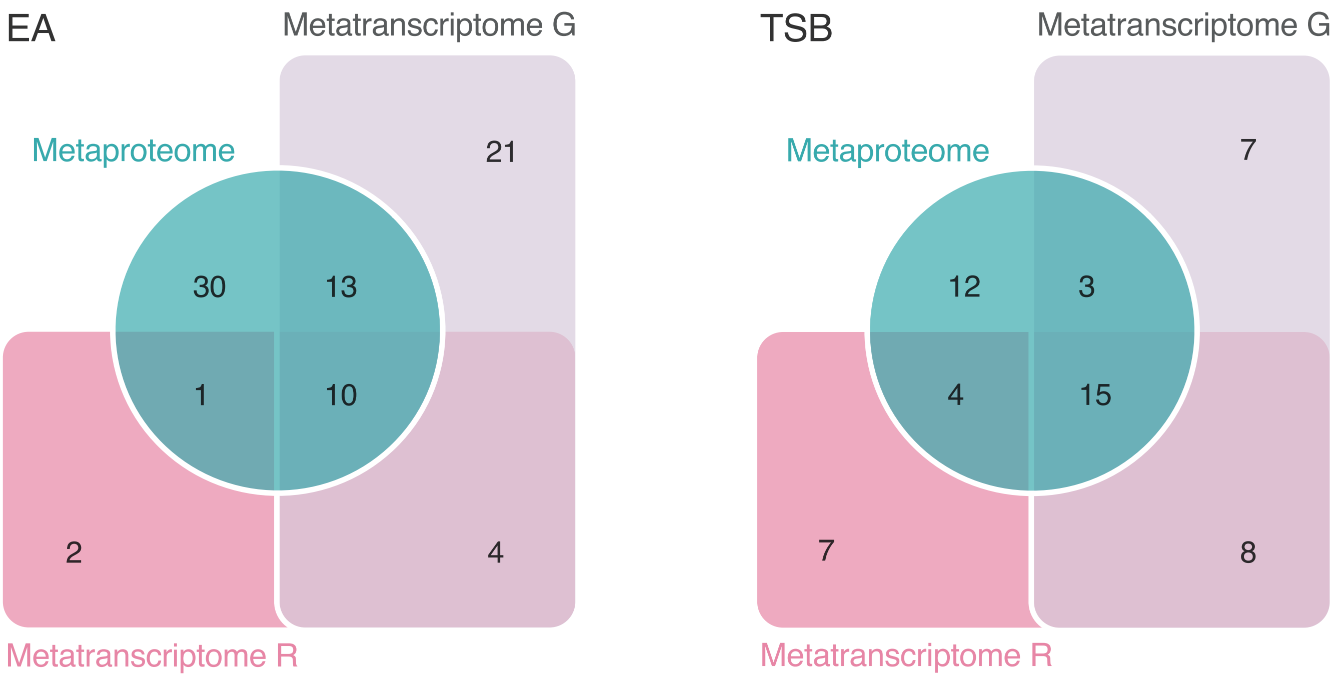


**Figure S15. Edwards-Venn diagrams of *ars* genes with increased expression in +As condition EA and TSB relative to no-As control cultures.** Number of *ars* genes encoded in metagenomes, with increased expression in metatranscriptomes R and G or/and metaproteomes from +As condition EA culture (left panel) and +As condition TSB (right panel) cultures.
