## Supplemental Material&Methods and Results for "Meta-omics-aided isolation of an elusive anaerobic arsenic-methylating soil bacterium"

### Supplementary Information (SI) for the research article: Meta-omics-aided isolation of an elusive anaerobic arsenic-methylating soil bacterium

Karen Viacava[1,2], Jiangtao Qiao[1], Andrew Janowczyk[3], Suresh Poudel[4], Nicolas Jacquemin[5], Karin Lederballe Meibom[1], Him K. Shrestha[4,6], Matthew C. Reid[7], Robert L. Hettich[4], Rizlan Bernier-Latmani[1*].

[1]Environmental Microbiology Laboratory, School of Architecture, Civil and Environmental Engineering, École Polytechnique Fédérale de Lausanne, Lausanne, Switzerland.
[2]Soil Science Group, Institute of Geography, University of Bern, Bern, Switzerland.
[3]Bioinformatics Core Facility, Swiss Institute of Bioinformatics, Lausanne, Switzerland.
[4]BioSciences Division, Oak Ridge National Laboratory, Oak Ridge, TN, United States.
[5]Translational Bioinformatics and Statistics (TBS), Department of Oncology, Université de Lausanne, Lausanne, Switzerland.
[6]Genome Science and Technology Graduate School, University of Tennessee, Knoxville, TN, United States.
[7]School of Civil and Environmental Engineering, Cornell University, Ithaca, NY, United States.

#### SI Materials and Methods

*DNA sequencing and metagenomic analysis*

DNA was extracted from the pellet (10 min, 4,500 g) of 4 ml of culture using the DNeasy® Power® Soil Kit (Qiagen, Hilden, Germany) with the modification of homogenizing with a Precellys 24 Tissue Homogenizer (Bertin Instruments, Montigny-le-Bretonneux, France) (6,500 rpm for 10 s, repeated 3x with 10s pause intervals). Metagenomic sequencing was performed by the Genomics Platform of the University of Geneva, Switzerland (iGE3) on an Illumina HiSeq 4000 (Illumina, San Diego, CA, US). Libraries were multiplexed and prepared using 100-base reads with paired ends according to the Nextera™ DNA Flex Library Preparation Kit protocol (Illumina). The quality of sequence reads was assessed with FastQC [1] and duplicated reads eliminated by FastUniq [2]. Reads from all biological replicates within the same experimental condition were assembled into contiguous sequences (contigs) using MegaHit [3]. The contig abundance was determined by aligning the sequencing reads from each biological replicate back to the assembled contigs using Kallisto [4, 5]. The abundance for each gene was considered equivalent to the abundance of the contig in which it was encoded. Gene abundance is reported as 'transcripts per million' (TPM), referred as TPM-DNA when used for gene abundance. TPM is a measure of gene abundance where reads are normalized for gene length and number of overall reads sequencing depth [6]. Prodigal was used for the prediction of protein-coding genes [7] generating protein sequence libraries for each culture (EA, TSB) and condition (no-As control, +As condition). The annotation server GhostKOALA [8] was used to assign a KEGG Orthology (KO) database number to each protein-coding gene to identify its encoded function and taxonomic category. The small subunit (SSU; 16S) rRNA sequences were identified directly in the contigs and their taxonomy assigned by Metaxa2 [9]. The relative abundance of the 16S SSU rRNA sequences identified in each of the four metagenomes was quantified using the contig abundance calculated by Kallisto. Contigs with length >2,000 bp were clustered into bins based on composition and coverage using CONCOCT [10], MetaBAT2 [11] and MaxBin 2.0 [12]. The final bin set was obtained by using the Bin_refinement module from MetaWRAP [13]. Completeness, contamination, strain heterogeneity and % community in contigs for each bin were calculated using CheckM [14]. Matching bins between the no-As and +As metagenomes, and between the EA +As and TSB +As metagenomes were found by pairwise comparison of the predicted genomes by using dRep [15]. Bins with an average nucleotide identity >95% were considered as identical genomes.

*RNA sequencing and metatranscriptomic analysis*

Each culture (5 ml) was harvested at mid-exponential phase for metatranscriptomic analysis. The cells were lysed and the RNA purified using the RNeasy Mini Kit following the manufacturer's instructions (RNAprotect® Bacteria, Qiagen). The purified RNA was DNase-I treated (Promega, Madison, WI, US) (1h, 37 °C) and cleaned using the RNeasy Mini Kit a second time. Ribosomal RNA (rRNA) depletion (kit QIAseq FastSelect −5S/16S/23S, Qiagen), library preparation using single-end 100 bases reads (TrueSeq Stranded mRNA, Illumina) and RNA sequencing (on an Illumina HiSeq 4000) were performed by the iGE3 Platform. Reads were quality-assessed by FastQC, trimmed by Trimmomatic [16], post-sequencing rRNA-depleted by SortMeRNA [17] and aligned to their corresponding protein sequence library by Bowtie2 [18]. The program featureCounts [19] was employed to count the number of RNA reads aligned to the Prodigal-predicted protein-coding genes. The raw counts were used to calculate the TPM, referred as TPM-RNA when employed for transcript abundance. Finally, to assess RNA expression changes in the +As condition relative to the no-As condition, a differential abundance analysis was performed using the DESeq2 package [20] using the protein sequence libraries from the +As condition to align the RNA reads. A gene was considered to have a significant difference in transcription when the absolute $\log_2$ fold change was ≥1 (i.e., 0.5 ≥fold change ≥2) and the adjusted q-value ≤0.05.

*Metaproteome characterization and metaproteomic analysis*

The metaproteome analysis was performed at Oak Ridge National Laboratory (Oak Ridge, TN, US). Biomass pellets from 100 ml of culture were washed with 100 mM $NH_4HCO_3$ buffer

(ABC) (pH 8.0), re-suspended in lysis buffer (4% sodium dodecyl sulfate, 100 mM Tris-HCl, pH 8.0) and disrupted by bead-beating. Lysate proteins were reduced with 5 mM dithiothreitol (30 min, 37 °C), alkylated with 15 mM iodoacetamide (30 min in the dark, room temperature) and isolated by a chloroform-methanol extraction. Extracted proteins were solubilized in 4% sodium deoxycholate (SDC) in ABC and the concentration estimated with a Nanodrop (Thermo Fisher Scientific, Waltham, MA, US). Sequencing-grade trypsin (Promega) at a 1:75 enzyme:protein ratio (w/w) was used to digest the proteins and formic acid (1% final concentration) was used to precipitate the SDC and collect tryptic peptides (peptide products of trypsin digestion). Aliquots of 12 μg of peptides were analyzed by 2D LC-MS/MS consisting of a Vanquish UHPLC connected to a Q Exactive Plus MS (Thermo Fisher Scientific). Spectral data were collected using MudPIT (multidimensional protein identification technology) as described previously [21, 22]. Peptides were separated in three steps (35, 100, and 500 mM ammonium acetate eluent) with organic gradients after each step. Eluted peptides were measured and sequenced by data-dependent acquisition using previously described parameters [23].

Protein databases were created for the +As experimental condition (i.e., EA +As and TSB +As) from the corresponding protein sequence libraries generated by Prodigal. The MS/MS spectra raw files were processed in Proteome Discoverer version 2.4 (Thermo Fisher Scientific) with MS Amanda 2.0 [24] and Percolator [25]. Spectral data were searched against the protein database of the corresponding culture (i.e., EA or TSB). The following parameters were used in the search algorithm MS-Amanda 2.0 to derive tryptic peptides: MS1 tolerance = 5 ppm; MS2 tolerance = 0.02 Da; missed cleavages = 2; carbamidomethyl (C, + 57.021 Da) as static modification; and oxidation (M, + 15.995 Da) as dynamic modifications. The false discovery rate (FDR) threshold was set to 1% for strict FDR and 5% for relaxed FDR at the peptide-spectrum matched (PSM), peptide, and protein levels. FDR-controlled peptides were then quantified according to the chromatographic area-under-the-curve and mapped to their respective proteins. Areas were summed to estimate protein-level abundance.

For differential abundance analysis of proteins, the spectral data from the no-As control (i.e., EA no As and TSB no As) were searched against the EA +As and TSB +As protein databases, respectively. All the above-described parameters were maintained. The proteins with at least 1 peptide evidence were exported from Proteome Discoverer. Protein data matrix from EA no As and EA +As were merged and TSB no As and TSB +As were merged. Protein abundance values were $\log_2$ transformed, LOESS-normalized between the biological replicates and mean-centered across all the conditions using InfernoRDN software [26]. Stochastic sampling of the proteins was filtered by removing the proteins that do not have abundance value present in at least two of the biological triplicates in at least one condition (no-As control or +As condition). Remaining missing data were imputed by random numbers drawn from a normal distribution (width = 0.3 and downshift = 2.8 using the Perseus software http://www.perseus-framework.org) [27]. The differentially abundant proteins were identified by Student t-test method with adjusted q-value ≤0.05. Proteins were further filtered using the absolute $\log_2$ fold change ≥1.

*Arsenic methylation assay for* Paraclostridium *sp. EML*

The As methylation assay consisted of a time-course experiment. RCB was brought to a boil, then cooled down to room temperature under a gas flow (100% $N_2$), and dispensed into 200-ml serum bottles (100 ml of medium per bottle) under the same gas atmosphere. The medium was amended with 25 μM As(III) as $NaAsO_2$, and a no-As(III) control was included (three replicates per condition). Bottles were inoculated with 1% inoculum (v/v) of a pre-grown exponential phase culture. Samples were obtained for soluble and volatile As as well as for microbial growth as previously described for anaerobic cultures [28]. Growth was quantified using $OD_{600}$. Arsenic bound to the biomass was assessed, as described in [29], by collecting a pellet of 1 ml of culture (13,000 g, 10 min), resuspending in 1 ml lysis buffer (0.1% triton X-100, 0.1% SDS, 10 mM EDTA, and 1 mM Tris−HCl), boiling 95 °C for 18 min with vortexing every 3 min and diluting 10x in 1%$HNO_3$ prior to analysis for total As and As species.

Arsenic speciation was determined in the step-gradient elution mode with an As Spec anion exchange fast column (50 mm x 4.0 mm, PrinCen, Guangzhou, China) (pump: 1.2 ml $min^{-1}$, injection volume: 10 μl, autosampler: 4°C and column compartment: 20°C). The mobile phase consisted of eluent A (0.23 ml 69% $HNO_3$ + 1.8 ml 28% ammonium) and eluent B (2.32 ml 69% $HNO_3$ + 4.6 ml 28% ammonium). The gradient program was set for 0-80 s 100% eluent A, 81-260 s 100% eluent B, 261-300 s 100% eluent A. Five arsenic standards were prepared, including TMAsO as trimethyl arsine oxide (Argus Chemicals Srl., Italy), DMAs(V) as sodium dimethylarsinate (ABCR, Germany), MMAs(V) as monomethylarsonic acid (Chemservice, PA, USA), and As(V) as $Na_2HAsO_4 \cdot 7H_2O$ (Sigma-Aldrich, MO, USA). Total As concentrations were measured using the same ICP-MS instruments in stand-alone mode [28]. Instrument settings in Table S1.

#### SI Results

*Arsenic methylation by soil-derived microbiomes*

The added initial As(III) concentrations, or those after spiking in the second set-up, were 28.3± 2.7 μM and 24.4± 5.8 μM for the EA and TSB experiments, respectively (Figures S1-B and S2-B). A decrease in the total initial soluble As was observed in most cases and could be due either to volatilization or intracellular accumulation of As (not assessed). Soluble methylarsenicals were found in all experiments, mainly mono- and dimethylated As (panels C and D from Figures S1 and S2). Arsenic species were oxidized prior to analysis [28]. Thus, As-speciation analysis discriminated between inorganic As and mono-, di-, or trimethylated arsenicals but did not allow the identification of the redox state of the methylarsenicals. Therefore, even when DMAs(V) and MMAs(V) were measured, they could have corresponded to trivalent species prior to oxidation.

The highest methylation efficiencies were achieved 45 h after inoculation: 27.7% of the initial As(III) for the EA culture, and 19.5% for the TSB culture. Previously, the EA culture was reported to convert 63% of the initial iAs 100 h after inoculation [29]. The variation in methylation efficiency between the previous work is likely due to the difference in sampling time. In the case of TSB experiment, the higher TSB methylation efficiency observed in the metagenome culture may be due to the lower initial concentration (15.1± 2.7 μM), as lower As concentrations have been observed to lead to higher As methylation efficiencies [28].

*Abundance changes in microbiome composition*

In the no-As control EA culture, Firmicutes (76.0 ±0.4%) was the most abundant phylum, particularly members of the order Clostridiales (49.6 ±1.6%) and Lactobacillales (16.9 ±1.2%) (Table S14). When the community was grown in the presence of arsenic, the taxa belonging to Clostridiales and Desulfovibrionales did not present statistically significant changes in abundance (Table S14). In contrast, significant changes were observed in the abundance of taxa from the order Lactobacillales, which decreased by two thirds, from 16.9 ±1.2% (no-As control) to 5.0 ±0.6% (+As condition), from the order Bacteroidales, which doubled going from 8.4 ±0.5% to 16.9 ±0.4%. Additionally, Selenomonadales and Acidaminococcales exhibited a slight but statistically significant increase from 4.2 ±0.5% to 5.1 ±0.3% and from 2.9 ±0.0% to 4.0 ±0.5%, respectively.

The order Enterobacterales dominated the microbiome in the TSB no-As control representing 65.5 ±6.4% of the community, followed by orders from the phylum Firmicutes: Clostridiales (24.7 ±4.7%), Lactobacillales (2.3 ±0.1%) and Bacillales (1.1 ±0.4%) (Table S14). The TSB community was altered upon exposure to As and exhibited a statistically significant increase in the order Lactobacillales (to 6.8 ±1.96%) and Bacteroidales (to 2.33 ±0.57%).

At the genus level (Figure 1, Table S15), the order Clostridiales included primarily contributions from members of the genera *Clostridium* and *Oscillibacter* for both EA and TSB. In addition, there was a contribution from organisms for which there is no attribution at the genus level (*Incertae Sedis*) which was especially large in the EA culture. In both cultures, Bacteroidales consisted entirely of the *Bacteroides* genus and Lactobacillales of the *Enterococcus* genus. Additionally, for EA, the orders Desulfovibrionales and Bacillales each comprised one genus, *Desulfovibrio* and *Bacillus*, respectively. In the TSB culture, the most abundant genera were *Citrobacter* and *Enterobacter*, in the absence and presence of arsenic respectively, both from the dominant Enterobacterales order.

*Abundance changes in MAGs*

The relative abundance of each MAG in the community was extracted using the *Profile* command in CheckM and reported as % Community. This percentage is calculated as the ratio of the number of reads mapped to the contigs in each MAG and the total number of reads mapped to all contigs (including the unbinned contigs), and adjusted for the size of the MAG (assuming an average genome size for the unbinned fraction). Based on % Community, approximately 77.6% and 38.0% of the EA and TSB+As microbiomes, respectively, are represented in the retained MAGs (Table 1). The most abundant MAGs, >5% relative abundance, in the EA community are: Clostridiales MAG 5 (25.21 ±0.23%), Deltaproteobacteria MAG 14 (12.71 ±0.49%), Clostridiales MAG 6 (12.68 ±0.68%), Bacteroidales MAG 1 (7.26 ±0.32%) and Clostridiales MAG 3 (5.73 ±0.24%); and, in the TSB community are: Bacteroidales MAG 1 (8.47 ±1.89%) and *Enterobacteriaceae* MAG 16 (6.77 ±0.35%).

The changes in % Community between the +As condition and the no-As control were significant in 14 of 17 the EA MAGs for which a matching MAG was found (Figure S10, upper panel). Some of the changes in the EA MAG abundances agree with the ones observed at the order level in the 16SS rRNA OTUs, e.g., the Bacteroidales MAG 1, and Selenomonadales MAG 19 increased as did their corresponding orders (Figure 1 and Table S13). However, the increment in both Deltaproteobacteria EA MAGs was not reflected in the 16SS rRNA OTUs.

Contrary to EA, most of TSB MAGs did not present statistically-significant abundance changes (Figure S10, lower panel) between the no-As control and +As condition. However, as in the case of EA, some of the changes observed agree with the 16SS rRNA OTU changes like the significant decrease in the Enterobacteriaceae MAG 16 as in the Enterobacterales order (Figure 1 and Table S14). Finally, most of the TSB MAGs with an equivalent MAG pair in the EA MAGs (Table S9), presented the same increasing or decreasing behavior in their abundance, except from the Bacteroidales and Selenomonadales MAGs that increased their community abundance in the EA +As culture compared to the no-As control while decreasing their abundance in the TSB +As culture compared to the no-As control.

*Active metabolic pathways in MAGs*

The presence, transcription and translation of genes encoding key enzymes from various metabolic pathways were assessed for each EA and TSB MAG (Figures S5 and S9). As described in the manuscript, a gene was considered as present in the MAG if DNA reads represented >5 TPM-DNA, as transcribed if RNA reads were >5 TPM-RNA and as translated if protein abundance could be calculated from the detected peptides, in at least two of the three biological replicates for each case. Individual values of gene, transcript and protein abundance levels for each MAG, biological replicate and enzyme are available in SI Tables S16 and S17.

The Deltaproteobacteria MAGs expressed the pathway for dissimilatory sulfate reduction (DSR) in both cultures; the dissimilatory reduction of nitrate to ammonia (DNRA) in the EA culture, presumably due to the presence of nitrate in the medium; and enzymes for acetate (AckA), ethanol (YiaY) and butanediol fermentation (as indicated by the expression of the acetolactate synthase, synthesizing a butanediol intermediary).

All members from the *Enterobactericeae* family are able to couple nitrate reduction to glucose fermentation[30]. The *Enterobactericeae* MAGs from TSB, expressed the enzymes needed for denitrification and mixed-acid fermentation: formate (PfD), acetate (AckA), ethanol (ADH5, and ADHE) and butanediol. Lactobacillales MAGs in both enrichments presented the same fermentative metabolisms as Enterobacterales plus lactate (LDH) and, in one of them, the amino-acid fermentation (Stickland fermentation).

Finally, in both cultures, the MAGs belonging to the phylum Firmicutes (Clostridiales, *Clostridium*, Lactobacillales, Selenomonadales and Firmicutes MAGs) displayed various mixed acid fermentations, acetone-butanol-ethanol (ABE) fermentation and amino-acid fermentation. Key enzymes from the acetyl-CoA (Wood-Ljungdahl) pathway, specially the formate-tetrahydrofolate (THF) ligase (Fhs), were expressed in all of the MAGs belonging to the phylum Firmicutes with the exception of Selenomonadales MAGs. Given the highly nutritious broth used for the growth of the communities, i.e., high carbon availability, the transcription and translation of *fhs* could be an indication of an active reverse Wood-Ljungdahl pathway. This reversal of the Wood-Ljungdahl pathway consists in the oxidation of acetate to hydrogen and carbon dioxide and has been shown to occur in anaerobic environments [31].

*Metatranscriptomics and metaproteomics differential analyses in metagenomic libraries*

Differential analysis of the metatranscriptomes shows that the presence of As impacted the transcription of a higher percentage of genes in the TSB culture than in the EA culture (Figure S11 and Table S18). For metatranscriptomes R, while in TSB culture the percentage of genes with increased or decreased transcription was almost the same, for EA culture the percentage of genes with increased transcription was greater that of the genes with decrease transcription. Conversely, in metatranscriptome G the percentage of genes with increased transcription was half of the genes with decreased transcription for TSB culture but remained almost the same for EA culture. In the case of the metaproteomes, the percentage of genes with increased expression was greater than the genes with decreased expression for EA and TSB cultures (Figure S12). The genes with differentially abundant RNA transcripts and proteins are enlisted in Tables S19 and S20.

*Identification of arsenic resistance genes*

In addition to GhostKOALA, the protein sequence libraries from EA and TSB of the +As condition were further annotated using the EggNOG server [32] in order to assign a KO number and an orthologous group (OG) to each protein-coding gene. Using the annotation from both servers, the libraries were queried for arsenic resistance (*ars*) genes (Table S21). To verify the annotation of the putative *ars* genes, the amino acid sequences of the encoded proteins were aligned, using BLAST®, against representative arsenic-resistance proteins (Table S22) selected based on previous studies [33, 34]. Alignments were considered accurate only when the E-value <0.01. The BLAST® annotation was assigned according to the annotation of either the top or the top three reference protein alignments. Further verification of the annotation was done by searching the amino acid sequences against Hidden Markov Model (HMM) profiles from the reference sequences using *hmmscan* in HMMER v3.1b2 package [35] with an E-value <0.01. One gene annotated by the databases as *arsB,* from EA +As library, was reassigned to *acr3* based on BLAST® and HMMER results (protein id k119_31951_258, Table S10).

*Increased expression of arsenic resistance genes*

In 23 of the 41 MAGs from the +As cultures, at least one *ars* gene exhibited increased transcription or protein expression relative to the no-As control (Figures S13 and S14). Only four *ars* genes, two *arsR2* in EA, an *arsC1* and an *arsP* in TSB showed a decrease, all four in the metaproteome. With the exception of Clostridiales MAGs 9 and 10 in TSB, all MAGs contained at least one *ars* gene (Figures S13 and S14).

A similar number of *ars* genes had increased transcription, relative to the no-As control, in both metatranscriptomes (R and G) for the TSB +As culture. In contrast, for the EA +As culture, threefold as many genes had increased RNA reads, relative to the no-As control, in metatranscriptome G as compared to metatranscriptome R (bold numbers in Figure 2). There was a significant overlap of *ars* genes, 44% in EA and 64% in TSB, that were detected as transcripts and proteins (Figure S15).

The expression as mRNA transcripts and/or proteins of *arsA*, *arsD*, *arsC*, *acr3*, and *arsB* in 21 of the 23 MAGs with expressed *ars* genes, underscores the predominance of intracellular As(III) efflux as a detoxification strategy in the microbiomes. The vast majority of As(III) efflux genes correspond to *acr3*, the most common membrane-transporter-encoding gene in *ars* operons [36]. The *arsB* gene was only expressed in the *Enterobacteriaceae* MAGs (TSB MAGs 15 and 16) in TSB (Figure S14).

The genes reported as unbinned consist of *ars* genes in contigs that could not be clustered in a MAG or that belonged to unretained, low-quality bins.

#### References


1. Andrews S. FastQC v0.11.9. https://www.bioinformatics.babraham.ac.uk/projects/fastqc/. .
2. Xu H, Luo X, Qian J, Pang X, Song J, Qian G, et al. FastUniq: A Fast De Novo Duplicates Removal Tool for Paired Short Reads. *PLoS One* 2012; **7**: e52249.
3. Li D, Liu CM, Luo R, Sadakane K, Lam TW. MEGAHIT: An ultra-fast single-node solution for large and complex metagenomics assembly via succinct de Bruijn graph. *Bioinformatics* 2015; **31**: 1674–1676.
4. Bray NL, Pimentel H, Melsted P, Pachter L. Near-optimal probabilistic RNA-seq quantification. *Nat Biotechnol* 2016; **34**: 525–527.
5. Schaeffer L, Pimentel H, Bray N, Melsted P, Pachter L. Pseudoalignment for metagenomic read assignment. *Bioinformatics* 2017; **33**: 2082–2088.
6. Wagner GP, Kin K, Lynch VJ. Measurement of mRNA abundance using RNA-seq data: RPKM measure is inconsistent among samples. *Theory Biosci* 2012; **131**: 281–285.
7. Hyatt D, Chen GL, LoCascio PF, Land ML, Larimer FW, Hauser LJ. Prodigal: Prokaryotic gene recognition and translation initiation site identification. *BMC Bioinformatics* 2010; **11**: 119.
8. Kanehisa M, Sato Y, Morishima K. BlastKOALA and GhostKOALA: KEGG Tools for Functional Characterization of Genome and Metagenome Sequences. *J Mol Biol* 2016; **428**: 726–731.
9. Bengtsson-Palme J, Hartmann M, Eriksson KM, Pal C, Thorell K, Larsson DGJ, et al. Metaxa2: Improved identification and taxonomic classification of small and large subunit rRNA in metagenomic data. *Mol Ecol Resour* 2015; **15**: 1403–1414.
10. Alneberg J, Bjarnason BS, De Bruijn I, Schirmer M, Quick J, Ijaz UZ, et al. Binning metagenomic contigs by coverage and composition. *Nat Methods* 2014; **11**: 1144–1146.

11. Kang DD, Li F, Kirton E, Thomas A, Egan R, An H, et al. MetaBAT 2: An adaptive binning algorithm for robust and efficient genome reconstruction from metagenome assemblies. *PeerJ* 2019; e7359.

12. Wu YW, Simmons BA, Singer SW. MaxBin 2.0: An automated binning algorithm to recover genomes from multiple metagenomic datasets. *Bioinformatics* 2016; **32**: 605–607.

13. Uritskiy G V, Diruggiero J, Taylor J. MetaWRAP - A flexible pipeline for genome-resolved metagenomic data analysis 08 Information and Computing Sciences 0803 Computer Software 08 Information and Computing Sciences 0806 Information Systems. *Microbiome* 2018; **6**.

14. Parks DH, Imelfort M, Skennerton CT, Hugenholtz P, Tyson GW. CheckM: Assessing the quality of microbial genomes recovered from isolates, single cells, and metagenomes. *Genome Res* 2015; **25**: 1043–1055.

15. Olm MR, Brown CT, Brooks B, Banfield JF. DRep: A tool for fast and accurate genomic comparisons that enables improved genome recovery from metagenomes through de-replication. *ISME J* 2017; **11**: 2864–2868.

16. Bolger AM, Lohse M, Usadel B. Trimmomatic: A flexible trimmer for Illumina sequence data. *Bioinformatics* 2014; **30**: 2114–2120.

17. Kopylova E, Noé L, Lè Ne Touzet H. SortMeRNA: fast and accurate filtering of ribosomal RNAs in metatranscriptomic data. 2012; **28**: 3211–3217.

18. Langmead B, Salzberg SL. Fast gapped-read alignment with Bowtie 2. *Nat Methods* 2012; **9**: 357–359.

19. Liao Y, Smyth GK, Shi W. Sequence analysis featureCounts: an efficient general purpose program for assigning sequence reads to genomic features. 2014; **30**: 923–930.

20. Love MI, Huber W, Anders S. Moderated estimation of fold change and dispersion for RNA-seq data with DESeq2. *Genome Biol* 2014; **15**: 550.

21. Washburn MP, Wolters D, Yates JR. Large-scale analysis of the yeast proteome by

multidimensional protein identification technology. *Nat Biotechnol* 2001; **19**: 242–247.

22. McDonald WH, Ohi R, Miyamoto DT, Mitchison TJ, Yates JR. Comparison of three directly coupled HPLC MS/MS strategies for identification of proteins from complex mixtures: Single-dimension LC-MS/MS, 2-phase MudPIT, and 3-phase MudPIT. *Int J Mass Spectrom* 2002; **219**: 245–251.

23. Clarkson SM, Giannone RJ, Kridelbaugh DM, Elkins JG, Guss AM, Michenera JK. Construction and optimization of a heterologous pathway for protocatechuate catabolism in escherichia coli enables bioconversion of model aromatic compounds. *Appl Environ Microbiol* 2017; **83**.

24. Dorfer V, Pichler P, Stranzl T, Stadlmann J, Taus T, Winkler S, et al. MS Amanda, a universal identification algorithm optimized for high accuracy tandem mass spectra. *J Proteome Res* 2014; **13**: 3679–3684.

25. Käll L, Canterbury JD, Weston J, Noble WS, MacCoss MJ. Semi-supervised learning for peptide identification from shotgun proteomics datasets. *Nat Methods* 2007; **4**: 923–925.

26. Polpitiya AD, Qian WJ, Jaitly N, Petyuk VA, Adkins JN, Camp DG, et al. DAnTE: A statistical tool for quantitative analysis of -omics data. *Bioinformatics* 2008; **24**: 1556–1558.

27. Tyanova S, Temu T, Sinitcyn P, Carlson A, Hein MY, Geiger T, et al. The Perseus computational platform for comprehensive analysis of (prote)omics data. *Nat Methods* . 2016. Nature Publishing Group. , **13**: 731–740

28. Viacava K, Meibom KL, Ortega D, Dyer S, Gelb A, Falquet L, et al. Variability in Arsenic Methylation Efficiency across Aerobic and Anaerobic Microorganisms. *Environ Sci Technol* 2020; **54**: 14343–14351.

29. Reid MC, Maillard J, Bagnoud A, Falquet L, Le Vo P, Bernier-Latmani R. Arsenic Methylation Dynamics in a Rice Paddy Soil Anaerobic Enrichment Culture. *Environ Sci Technol* 2017; **51**: 10546–10554.

30. Halkman HBD, Halkman AK. Indicator Organisms. *Encyclopedia of Food*

*Microbiology: Second Edition*. 2014. Elsevier Inc., pp 358–363.

31. Müller B, Sun L, Schnürer A. First insights into the syntrophic acetate-oxidizing bacteria - a genetic study. *Microbiologyopen* 2013; **2**: 35–53.

32. Huerta-Cepas J, Szklarczyk D, Heller D, Hernández-Plaza A, Forslund SK, Cook H, et al. EggNOG 5.0: A hierarchical, functionally and phylogenetically annotated orthology resource based on 5090 organisms and 2502 viruses. *Nucleic Acids Res* 2019; **47**: D309–D314.

33. Chen S-C, Sun G-X, Yan Y, Konstantinidis KT, Zhang S-Y, Deng Y, et al. The Great Oxidation Event expanded the genetic repertoire of arsenic metabolism and cycling. *Proc Natl Acad Sci* 2020; **117**: 10414–10421.

34. Rosen BP, Bhattacharjee H, Zhou T, Walmsley AR. Mechanism of the ArsA ATPase. *Biochim Biophys Acta - Biomembr* . 1999. , **1461**: 207–215

35. Johnson LS, Eddy SR, Portugaly E. Hidden Markov model speed heuristic and iterative HMM search procedure. *BMC Bioinformatics* 2010; **11**: 431.

36. Yang Y, Wu S, Lilley RM, Zhang R. The diversity of membrane transporters encoded in bacterial arsenic-resistance operons. *PeerJ* 2015; **3**: e943.
